# Tomato leaves under stress: A comparison of stress response to mild abiotic stress between a cultivated and a wild tomato species

**DOI:** 10.1101/2021.08.31.458379

**Authors:** Julia J. Reimer, Björn Thiele, Robin T. Biermann, Laura V. Junker-Frohn, Anika Wiese-Klinkenberg, Björn Usadel, Alexandra Wormit

**Author notes:** J.J. Reimer and B. Thiele contributed equally to this work. Corresponding author: Alexandra Wormit, RWTH Aachen University, Institute for Biology I, Worringer Weg 3, 52074 Aachen, Germany.

## Abstract

Tomato is one of the most produced crop plants on earth and growing in the fields and greenhouses all over the world. Breeding with known traits of wild species can enhance stress tolerance of cultivated crops. In this study, we investigated responses of the transcriptome as well as primary and secondary metabolites in leaves of a cultivated and a wild tomato to several abiotic stresses such as nitrogen deficiency, chilling or warmer temperatures, elevated light intensities and combinations thereof.

The wild species responded different to varied temperature conditions compared to the cultivated tomato. Nitrogen deficiency caused the strongest responses and induced in particular the secondary metabolism in both species but to much higher extent in the cultivated tomato. Our study supports the potential of a targeted induction of valuable secondary metabolites in green residues of horticultural production, that will otherwise only be composted after fruit harvest. In particular, the cultivated tomato showed a strong induction in the group of mono caffeoylquinic acids in response to nitrogen deficiency. In addition, the observed differences in stress responses between cultivated and wild tomato can lead to new breeding targets for better stress tolerance.

## 1 Introduction

Abiotic stresses and environmental changes have been shown to dramatically alter plant growth, metabolism and crop yield. Plants synthesize a variety of (plant) secondary metabolites (PSM) from primary metabolites often in response to various abiotic environmental signals (Løvdal et al., 2010; Selmar and Kleinwächter, 2013; Ramakrishna and Ravishankar, 2011; Junker-Frohn et al., 2019). PSM may act as signal molecules to elicit a stress response or scavenge excessive reactive oxygen species to minimize cellular damage (Wu et al., 2014).

Some major groups of PSM can be distinguished, such as phenylpropanoids or phenolic compounds, terpenoids, alkaloids, and sulfuric compounds (Crozier et al., 2007; Chomel et al., 2016). Phenolics are mainly represented by hydroxycinnamic acids, flavonoids, and anthocyanins, which collectively account for approximately 40 % of the organic carbon in the biosphere (Croteau et al., 2000). They can be subdivided in the large group of structural phenolics (like lignin, suberin and other structural polymers) and a minor fraction of non-structural phenolics with many functions in plants, including antioxidative activity (Grace and Logan, 2000; Grace, 2007). In addition, phenolic compounds are excellent oxygen radical scavengers (Bors et al., 1990; Grace, 2007). They can eliminate reactive oxygen intermediates without promoting further oxidative reactions (Grace, 2007).

Flavonoids are the most abundant polyphenols in plants and the human diet (Ock et al., 2007; Carvalho Lemos et al., 2019; Ruskovska et al., 2020). A wide variety of biological activities has been attributed to flavonoids, including protection against abiotic stresses like UV-B, cold stress, low nitrogen as well as defence against pathogens (Tohge et al., 2017), and tolerance to oxidative stress or drought (Nakabayashi et al., 2014). Accordingly, flavonoid biosynthesis can be stimulated by abiotic stresses like nutrient deficiency, elevated light intensities or cold temperatures (Løvdal et al., 2010; Lillo et al., 2008; Bénard et al., 2011). As an example, drought stress has been shown to induce phenolic compounds, di- and triterpenoids, alkaloids and other PSM up to 10 fold (Selmar and Kleinwächter, 2013). Flavonoids have beneficial properties for human health, including antioxidant, anti-retroviral, anti-hypertensive, anti-inflammatory, anti-aging and insulin-sensitizing activities (Yao et al., 2004; Grassi et al., 2015; Tohge and Fernie, 2017).

In field conditions, plants are often simultaneously exposed to various abiotic stresses, which have profound effects on crop yield and quality. Interactions of different stresses lead to different signalling events and transcriptional responses different from the single stresses (Rasmussen et al., 2013; Pandey et al., 2015). These alterations in gene expression lead to a specific regulation of the metabolism depending on the particular stress or stress combination. An increase in the amount of polyphenols and the antioxidant potential under stress conditions has been observed, indicating that the antioxidant mechanism is an essential strategy to guard plants against stress combinations but also in general (Løvdal et al., 2010; Pandey et al., 2015; Šamec et al., 2021). Thereby, the structure of the polyphenols and their glycosylation status will influence the antioxidative capacity (Šamec et al., 2021). However, most studies of abiotic stress responses focus on a single stressor, neglecting the confounding effect of multiple stresses on gene expression changes affecting crop yield and quality.

Tomatoes (*Solanum lycopersicum*) are one of the most widely cultivated fruits in the world with over 180 mio. tons produced in 161 countries in 2019 (http://faostat.fao.org/) and represent an important dietary source of bioactive compounds like antioxidants. However hundreds of years of tomato domestication has led to extensive loss of genetic variability (Ranc et al., 2008; Abewoy Fentik, 2017; Tamburino et al., 2020), indicated by only 5 % of the genetic and chemical diversity of wild relatives remaining in the cultivated tomato (Miller and Tanksley, 1990; Blanca et al., 2015; Fernie and Aharoni, 2019; Mata-Nicolás et al., 2020). One strategy of improving modern tomato varieties is to introduce quality traits from wild genetic resources that differ significantly in their phenotypic and agronomic traits, e.g. abiotic stress tolerance, or antioxidant content (Eshed and Zamir, 1995; Ofner et al., 2016; Mata-Nicolás et al., 2020).

The capacity to produce PSM in crop plants is often investigated for fruits or in medicinal plants (Isah, 2019), but rarely in remaining green residuals of agri- or horticulture. Horticultural production of tomatoes produces green residuals in the range of 33 kg per 100 kg produced tomato fruits as by-product (Boulard et al., 2011). Most of those residues will be composted, or (for a minor part) used for biorefinery (Wenger and Stern, 2019; De Buck et al., 2020). Previously, the specific induction of rutin production in tomato leaves under greenhouse conditions was shown by application of mild abiotic stresses (Junker-Frohn et al., 2019; Röhlen-Schmittgen et al., 2020). Based on those results, extraction of valuable PSM from leaves before composting was postulated, and will add a new aspect to the value-chain of horticulture.

The wild relative *Solanum pennellii* inhabits the coastal region of the Atacama Desert of Peru which is characterized by cool temperatures and hardly any precipitation. Correspondingly, *S. pennellii* shows adaptation to drought stress such as increased leaf thickness, lower stomatal frequency and a changed cuticle (Coneva et al., 2017; Fernandez-Moreno et al., 2017). In addition, the wild species also exhibit an increased salt stress tolerance partly due to up-regulated ROS scavenging systems (Ijaz et al., 2017). Besides, *S. pennellii* has a higher content of secondary metabolites in leaves and fruits (Schauer et al., 2004; Tohge et al., 2020). The availability of an introgression line population (Eshed and Zamir, 1995; Ofner et al., 2016) and genome sequence information (Bolger et al., 2014a; Schmidt et al., 2017) has assisted in identification of multiple QTL for different traits (e.g. fruit primary metabolite composition (Lin et al., 2014); volatile metabolites and antioxidants of fruits (Tieman et al., 2017); tolerance to abiotic and biotic stress; and secondary metabolite abundance (Alseekh et al., 2015) and candidate genes involved in drought stress tolerance (Fernandez-Moreno et al., 2017)). In particular, *S. pennellii* LA0716 is known to produce high amounts of flavonoids in dry seeds compared to other accessions of *S. pennellii* (Alseekh et al., 2020).

Due to its importance as genetic resource and experimental model, we utilized *S. pennellii* LA0716 in this study to advance our understanding of the underlying molecular mechanisms of abiotic stress response in tomato leaves to moderate temperature changes (chilling, warmth), nitrogen deficiency, and elevated light intensities as well as stress combinations thereof.

## 2 Materials and Methods

### 2.1 Plant cultivation

*Solanum lycopersicum* var. Lyterno (Rijk Zwaan Welver GmbH, Germany) and *Solanum pennellii* LA0716 (Tomato Genetics Resource Center, UC Davis, CA, US) were used to compare the effects of abiotic stress.

In brief, plants were grown from seeds in environmental chambers (Hühren Kälte-Klima-Elektrotechnik, Erkelenz, Germany), equipped with metal halide lamps (Philips, Hamburg, Germany), following the protocol described in Junker-Frohn et al. (2019), under controlled conditions of 22/18 °C during day and night, a relative humidity of 50 % and 200 µmol m^−2^ s^−1^ photons light intensity for 10 h per day.

Plants were sown in rock wool plugs (2×2×4 cm; Grodan, Roermond, The Netherlands), that were four times prewashed with deionized water. Seeds were placed around 0.5 cm below the surface of the rock wool plugs. A total number of 150 seeds per species were seeded. The plugs were watered with water for 16 days. 84 seedlings per species, which already developed the first true leaf, were then transferred to rock wool blocks (7.5×7.5×6.5 cm) (Grodan, Roermond, The Netherlands), that were three times prewashed with deionized water. Afterwards, the seedlings were fertilized with half-strength Hoagland solution for 14 days, followed by full-strength Hoagland solution (5 mM KNO_3_, 5 mM Ca(NO_3_)_2_, 2 mM MgSO_4_, 1 mM KH_2_PO_4_, 90 µM FeEDTA, plus micronutrients) for further 11 days.

### 2.2 Abiotic stress treatments and harvesting

The chosen abiotic conditions (nitrogen deficiency, chilling temperatures, elevated light intensities) were described previously (Løvdal et al., 2010; Junker-Frohn et al., 2019). In addition, a separate biological experiment was conducted with warmer temperature regimes, in equivalent walk-in chambers (for overview of experimental set up see also supplementary table 2). All plants in chambers were randomized once a week.

In brief, six weeks after germination, plants were stressed by nitrogen deficiency (N-), chilling temperatures (cold), warmer temperature regime (warm) or elevated light intensity (eL) and combinations thereof (N-cold, N-eL, eLcold and N-eLcold). Nitrogen deficiency was initiated by washing of the substrate and subsequent watering with nitrogen-free modified Hoagland solution (2.5 mM K_2_SO_4_, 5 mM CaCl_2_, 2 mM MgSO_4_, 1 mM KH_2_PO_4_, 90 µM FeEDTA, plus micronutrients). To remove most of the nitrogen, we washed the rockwool cubes three times with nitrogen-free medium.

Chilling temperatures were achieved by transferring plants to an identical environmental walk-in chamber set to 20/12 °C (day/night). Light intensity was doubled by elevating plants by 80 cm. Warmth stress (warm) was achieved by transferring plants to an identical environmental walk-in chamber set to 24/20 °C (day/night). An additional measurement of the canopy or leaf temperature was not performed. After one week of stress treatment, leaflets of the fourth leaf (counted from the tip) were sampled, immediately frozen in liquid nitrogen and stored at −80 °C. The stress treatments were performed in two independent experiments.

A total number of 68 tomato plants per species were harvested. At day 0 (d0), four control plants were harvested each. At day 4 (d4) and day seven (d7) after induction of the stress treatment four plants per stress and four control plants were harvested each.

Plant homogenization to a fine powder was done by grinding using liquid nitrogen, mortar and pistil. Afterwards, samples were stored at −80 °C.

### 2.3 Total RNA extraction and DNase-digestion

RNA was extracted by two different approaches: 1) for RNASeq: RNA was extracted using a column based kit from QIAGEN GmbH (Plant RNeasy, QIAGEN GmbH, Hilden, Germany) according to the manufacture’s protocol, 2) for cDNA synthesis and following quantitative real time PCR: a TRIZOL®-approach (Thermo Fisher Scientific Life Technologies GmbH, Darmstadt, Germany) was used according to the manufacture’s protocol. Both methods started with 50-100 mg ground tissue, and total RNA was eluted in 30-40 µL RNase-free water. The concentration and quality of the total RNA was determined by a Nanodrop measurement (Nanodrop 2000c, Thermo Fischer Scientific GmbH, Schwerte, Germany) and a native agarose gel electrophoresis.

To remove residual genomic DNA in the total RNA samples, a DNase digest was performed using Baseline-ZERO™ DNase (Epicentre, Madison, USA). 3.4 µL of the reaction buffer was added to 30 µL total RNA, in addition to 1 µL DNase (1 MBU). The digest was performed at 37 °C for 30 minutes. The DNase was inactivated by adding 4 µL stop buffer and an incubation at 70 °C for 10 minutes. Afterwards, total RNA was precipitated at −20 °C for 2 days using 100 µL isopropanol. The sample was washed twice using 70 % ethanol, air dried for 10 min at room temperature and resuspended in 30-40 µL RNase-free water.

### 2.4 RNASeq data handling and analysis

mRNA was enriched from total RNA samples, and subsequently analysed using an Illumina-platform (HiSeq) sequencing 2×75 bp paired-end reads.

Raw reads were trimmed using Trimmomatic (Bolger et al., 2014b). Reads were aligned to the respective genome or transcriptome (*S. pennellii* - Bolger et al. (2014a), *S. lycopersicum* built ITAG2.4 - Mueller et al. (2005); Fernandez-Pozo et al. (2015)) using hisat 2 (version 2.1.0) and salmon (Kim et al., 2019; Patro et al., 2015).

An artificial transcriptome was build using default settings of StringTie (Pertea et al., 2015, 2016) with trimmed reads of all analysed conditions, to investigate how similar the two species behave in a principal component analysis (PCA).

Read abundancy was then calculated by using salmon for each genome. Data analysis was performed using R 3.5.2 (R Core Team, 2019). Read abundancies were analysed using R-packages limma (Ritchie et al., 2015), edgeR (Robinson et al., 2009; McCarthy et al., 2012), and tximport (Soneson et al., 2016).

Overrepresentation analysis was carried out with PageMan (Usadel et al., 2006). Weighted cluster analysis was performed using the R package WGCNA (Langfelder and Horvath, 2008) (with default settings, softpower = 9, minModuleSize = 30, and mergeCutHeight = 0.25). During the analysis with WGCNA, genes with a low coefficient of variation among all sample types were discarded and the remaining genes were used for the analysis.

### 2.5 cDNA synthesis

Per 10 µL of RNA (1 µg total RNA) a total of 100 pmol oligo dT primers were added. The mix was heated to 70 °C for 5 minutes and cooled on ice afterwards. 8 µL of a mix consisting of 2 µL deoxyribonucleotide triphosphate (dNTPs, Promega, Madison, Wisconsin, US), 4 µL 5x RT-buffer, 0.5 µL reverse transcriptase (RT, Promega, Madison, Wisconsin, US) and RNase-free water were added. The reverse transcription was performed at 37 °C for 60 minutes and the reverse transcriptase was inactivated at 70 °C for 10 minutes. cDNA was stored at −20 °C until further analysis.

### 2.6 Quantitative realtime PCR

To analyse the relative gene expression of genes related to flavonoid biosynthesis in response to abiotic stress treatment a qRT-PCR was performed on a LightCycler® 480 II (Roche Diagnostics Deutschland GmbH, Mannheim, Germany) operated with LightCycler® 480 SW 1.5.1.62 software (Roche Diagnostics Deutschland GmbH, Mannheim, Germany). GoTaq® qPCR Master Mix (Promega GmbH, Mannheim, Germany) was used to carry out the qRT-PCR with oligomers as indicated in table 1. The reaction was performed in a total volume of 10 µL according to manufacturers instruction using white 384-well plates (Sarstedt AG & Co. KG, Nümbrecht, Germany) sealed with LightCycler® 480 Sealing Foil (Roche Diagnostics Deutschland GmbH, Mannheim, Germany). The PCR program was as followed: pre-denaturation: 180 s at 95 °C, 45 cycles of: 15 s at 98°C, followed by 20 s at 58 °C, and 20 s at 72 °C, and final elongation: 180 s at 72 °C. The melting curve was performed in the temperature range from 60 °C to 95 °C with an increase of 0.02 °C/s.

**Table 1:**
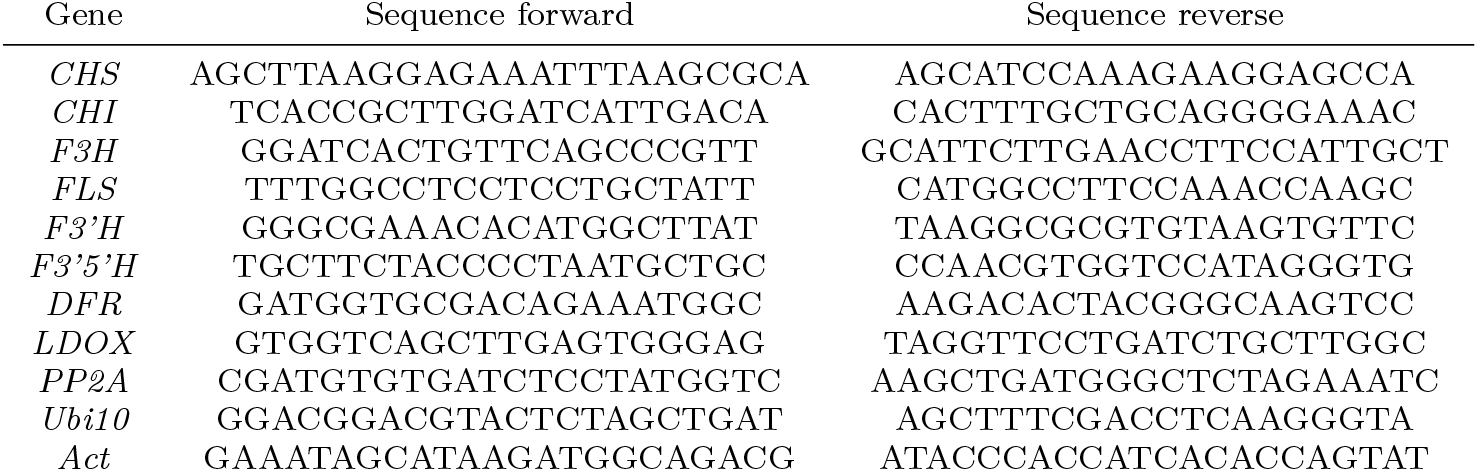
Used oligonucleotide primer pairs. Primer pairs were used with both species. The sequences are given in 5’-3’ direction. *CHS - Chalcone synthetase, CHI - Chalcone isomerase, F3H - Flavanone-3-hydroxylase, FLS - Flavonol synthase, F3’H - Flavonoid-3’-hydroxylase, F3’5’H - Flavonoid-3’,5’-hydroxylase, DFR - Dihydroflavonol-4-reductase, LDOX - Leucoanthocyanidin dioxygenase, PP2A - Proteinphosphatase 2, Ubi10 - Ubiquitin 10, Act - Actin*.

For analysis of the qRT-PCR data the ‘second derivative maximum method’ was performed with the LightCycler® 480 SW 1.5.1.62 software to calculate crossing point (Cp) values that are related to the maximal acceleration of fluorescence. Additionally, cDNA of all samples of one species were pooled to obtain a standard curve for determination of primer efficiency. The used dilutions in water were: 1:1, 1:10, 1:50, 1:100, 1:250 and 1:500.

Data obtained from qRT-PCR were analysed with REST-384© (Pfaffl, 2002). Significance was evaluated by a Pair Wise Fixed Reallocation Test©, performing 10.000 reallocations, implemented in REST 384©.

### 2.7 Extraction of metabolites

Fine-ground powder of the sample leaf were used to extract metabolites using 100 % methanol (final concentration 30 mg powder/ml methanol). The samples were mixed carefully and incubated for 4 h at room temperature in darkness with mixing every 90 min. Afterwards the samples were centrifuged at 4 °C for 15 minutes with 20800 rcf. The supernatants were filtered through a PVDF membrane with 0,2 µm pores into a new 2 mL reaction tube and stored at –80 °C till measurement.

For LC-FTICR-MS analysis sample extracts were directly used. For LC-MS/MS analysis 10 µL 50 µM quercetin-3-O glucuronide as internal standard (IStd) was added to 100 µL sample extract. Samples were further diluted till 500 µL total volume with 100 % methanol before analysis.

To analyse the metabolite composition we pooled samples of the same stress condition coming from one experiment and run those as a single sample. Two independent biological experiments were performed (see supplementary table 2).

### 2.8 Identification of secondary metabolites by LC-FTICR-MS and LC-MS/MS

Liquid chromatography-Fourier transform ion cyclotron resonance-mass mass spectrometry (LC-FTICR-MS) experiments were carried out using an Agilent 1200 series HPLC system consisting of a binary pump, autosampler, and column oven (Santa Clara, CA, USA). Secondary metabolites were separated on an Aqua 3 µm C18 column (150 x 2 mm, 3 µm particle size) equipped with a pre-column filter from Phenomenex. The mobile phase consisted of 1 mM aqueous ammonium acetate (A), and methanol + 1 mM ammonium acetate (B). Samples were separated at 40 °C and a flow rate of 0.3 mL/min using gradient elution: 30 % A isocratic for 1 min, linear gradient to 1 % A over 10 min, 1 % A isocratic for 20 min, linear gradient to 30 % A over 1 min and equilibration at 30 % A for 8 min (total run time: 40 min). The injection volume was 10 µL.

Mass spectrometry was performed using a hybrid linear ion trap-FTICR-mass spectrometer LTQ-FT Ultra (ThermoFisher Scientific, Bremen, Germany) equipped with a 7 T supra-conducting magnet. The electrospray ionization (ESI) source was operated in the positive as well as negative mode with a spray voltage of 3.70 and 2.80 kV, respectively. Nitrogen was employed as both sheath gas (8.0 arb) and auxiliary gas (0 arb). The transfer capillary temperature was set to 275 °C. Voltages for capillary and tube lens were set to 44 V and 175 V in ESI(+)-mode, and to −33 V and −135 V in ESI(-)-mode.

Mass spectra were recorded in a full scan from 100 to 1000 Da with a mass resolution of 100,000 at m/z 400 (full width at half maximum). The automatic gain control for providing a constant ion population in the ICR cell was set to 5E5 for the FTMS full scan mode. The maximum ion trap fill time was set to 10.0 ms and the maximum ICR cell fill time to 500 ms.

For ESI(+) mass spectra the accurate masses of quasi-molecular ions [M + H]^+^, [M + NH_4_]^+^ and [M + Na]^+^ were used for the calculation of chemical formulae with the Qual Browser in Xcalibur software version 2.0.7. For ESI(-) mass spectra it was the quasi-molecular ion [M - H]^-^.

The search algorithm for ESI(+) mass spectra contained the isotopes ^1^H, ^12^C, ^13^C, ^14^N, ^16^O and ^23^Na. For ESI(-) mass spectra ^14^N and ^23^Na were omitted. Each compound had to be represented by at least two mass peaks: the base peak and the peak(s) of the corresponding ^13^C-isotopologue(s). Search results were restricted to mass errors of ≤ 1.0 ppm for the ^12^C- and the corresponding ^13^C-isotopologue. Chemical formulae were used for searches of compounds against the ChemSpider and PubChem databases.

Liquid chromatography-tandem mass spectrometry (LC-MS/MS) was performed on a Waters ACQUITY^®^ UHPLC system (binary pump, autosampler) coupled to a Waters Xevo TQ-S^®^ triple-quadrupole mass spectrometer (Waters Technologies Corp., MA, USA). Separation of metabolites was achieved on a Nucleoshell RP18 column (100 x 4.6 mm, 2.7 µm; Macherey-Nagel, Germany). The column was equipped with a precolumn (Macherey–Nagel, Germany). The mobile phase were water (A) and acetonitrile (B) each containing 0.1 % formic acid, at a flow rate of 1.0 mL/min. The gradient program was as following: 98 % A to 65 % A within 10 min, to 0 % A within 0.1 min and holding for 2.4 min, back to 98 % A within 0.1 min and holding for 2.4 min. The injection volume was 10 µL.

Measurements of the metabolite extracts were done in the positive as well as negative ESI-mode. The capillary voltage was set to 2.5 kV in ESI(+) mode and −2.0 kV in ESI(-) mode. The desolvation temperature was 600 °C and the source temperature 150 °C. The desolvation gas flow was set to 1000 L/h and the cone gas flow at 150 L/h using nitrogen in both cases. Mass spectrometric detection was performed in the multiple reaction monitoring (MRM) mode with MRM transitions listed in supplementary table 2. Nitrogen was used as the collision gas at a flow of 0.15 mL/min. Metabolites were quantified by calculation of their peak area ratios A_Metabolite_/A_IStd_.

## 3 Results

### 3.1 Global response of the transcriptome to abiotic stresses is strongly influenced by nitrogen deficiency in both species

To investigate the effect of abiotic stress treatments in *S. lycopersicum* and *S. pennellii*, an RNASeq analysis from samples harvested at day 7 was performed. Reads from both species were mapped to a combined artificial transcriptome for a first overall comparison. A principal component analysis (PCA) showed that the highest level of variation is coming from the species (see supplementary figure 1 a), while the next level of variation is influenced by the stress treatments (see supplementary figure 1 b).

Afterwards, we mapped the reads to the respective reference genomes and performed the analysis for each species separate. In this context, the proportion of variance for both species was found to be similar, so that the main difference between samples can be explained by PC1 (44.97 % for *S. lycopersicum* and 37.57 % for *S. pennellii*) and PC2 (18.81 % for *S. lycopersicum* and 25.16 % for *S. pennellii*) (see figure 1 a and b). In addition, the PCA revealed that for both species samples coming from stress treatments including nitrogen deficiency (in any combination) grouped together (see orange dots in figure 1 a and b) and separated clearly from all other samples.

**Fig. 1:**
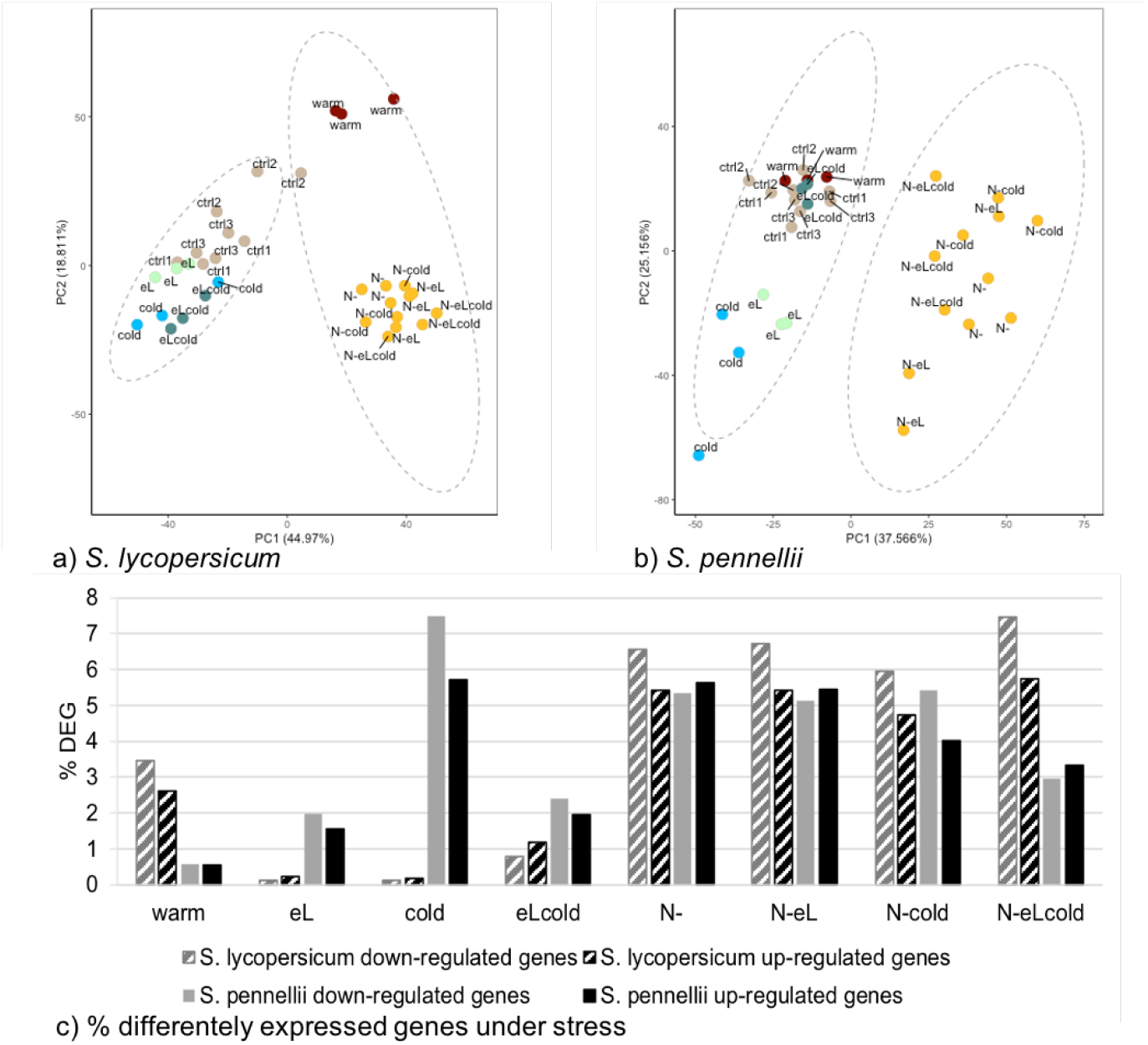
RNASeq analysis reveals different responses to temperature changes in the two tomato species. Total RNA was extracted from harvested material of day 7. (**a + b**) Principal component analysis (PCA) for *S. lycopersicum* (**a**) and *S. pennellii* (**b**) indicating responsivness to the stress treatments, shown by comparison of principal component 1 (PC1) versus principal component 2 (PC2). Control samples are indicated in brown, cold samples in blue, warm samples in red, elevated light samples in light green, and all samples comprising nitrogen deficiency in orange. (**c**) Percentage of differentaly expressed genes (DEG) in *S. lycopersicum* (striped columns) and *S. pennellii* (plain columns). Differentially expressed genes were obtained by global comparison of expression data for the stress treatment with the respective control sample. Percentage of down-regulated genes is shown in grey bars, and up-regulted genes in black bars, respectively. FDR<0.01

When performing a cluster analysis (see supplementary figure 2), we observed a similar pattern. Here all stresses harboring nitrogen deficiency form one single cluster for *S. lycopersicum* (see supplementary figure 2 a) but two distinct cluster for *S. pennellii* (see supplementary figure 2 b).

In *S. lycopersicum*, plants grown under warmer temperatures (warm) were more isolated from control samples (ctrl) (see figure 1 a, red dots vs. brown dots) compared to *S. pennellii* (see figure 1 b, red dots vs. brown dots). The single stresses with elevated light intensities (eL, green dots) or chilling temperatures (cold, blue dots) were more pronounced distinguishable from the control group for the wild *S. pennellii* than for the cultivated *S. lycopersicum*.

When checking the percentage of differentially expressed genes (DEG) by comparison with the respective control sample it becomes obvious that elevated light intensities and chilling temperature had the mildest effect in *S. lycopersicum* (eL and cold, see figure 1 c, and supplementary figure 3), and increased temperatures had the lowest impact on the DEG in *S. pennellii* (warm, see figure 1 c, and supplementary figure 3). Chilling temperatures (cold) gave only little DEG for the cultivated tomato *S. lycopersicum* while it had a strong effect in the wild relative *S. pennellii*. The opposite was found (although less pronounced) for warmer temperature conditions (warm). While the combination of elevated light regime and chilling temperatures has a strong effect for the cultivated tomato, less transcriptional changes could be observed in *S. pennellii* when compared to cold stress alone.

The percentage of DEG for up- and down-regulated genes was often similar in *S. pennellii*, while in *S. lycopersicum* the percentage of down-regulated genes was always slightly higher than the percentage of up-regulated genes. In all stresses harbouring nitrogen deficiency (N-), the amount of DEG was in between 3 % and 7 % for down- or up-regulated genes in both species (compare figure 1 c). We conclude from that observation that nitrogen deficiency had the highest impact in our approach and dominates the effects in combinations of abiotic stresses.

### 3.2 General differences in the transcriptomic response can be found for particular metabolic pathways

To get insights what metabolic functions were induced or repressed by the abiotic stress treatment, we first performed an over-/underrepresentation analysis (ORA) using the MapMan/PageMan ontology (Lohse et al., 2014; Usadel et al., 2009; Schwacke et al., 2019) in the subsets of up- or down-regulated genes, respectively.

During an ORA, the expression changes will be linked to potential biological responses. First, transcripts will be grouped in classes (or bins in case of PageMan), followed by an analysis if transcripts within the bin respond in a concerted way. In a simple ORA, a list of genes (here up- or down-regulated genes) is searched for biological functions or pathways that are enriched (by passing a threshold) compared to the complete list of functions/pathways, resulting in only overrepresentation indications. In addition, PageMan applies the same procedure for objects below the negative value of the threshold resulting in underrepresentation (negative value; depletion). In general, the overrepresentation of a category indicates a direct effect of the applied abiotic conditions (either induction for up-regulated genes or repression for down-regulated genes). In addition, the ORA will result in an underrepresentation, if the expression of the transcripts within a given bin is not affected by the applied conditions.

Our analysis revealed an overrepresentation (see table 2 shown as red dots) for the categories phytohormone action and secondary metabolism in up- and down-regulated genes of both species. Further functions show a more divers response to the applied abiotic stress conditions, often resulting in an overrepresentation of either up- or down-regulated genes, e. g. RNA biosynthesis, lipid metabolism, or external stimuli response.

**Table 2:**
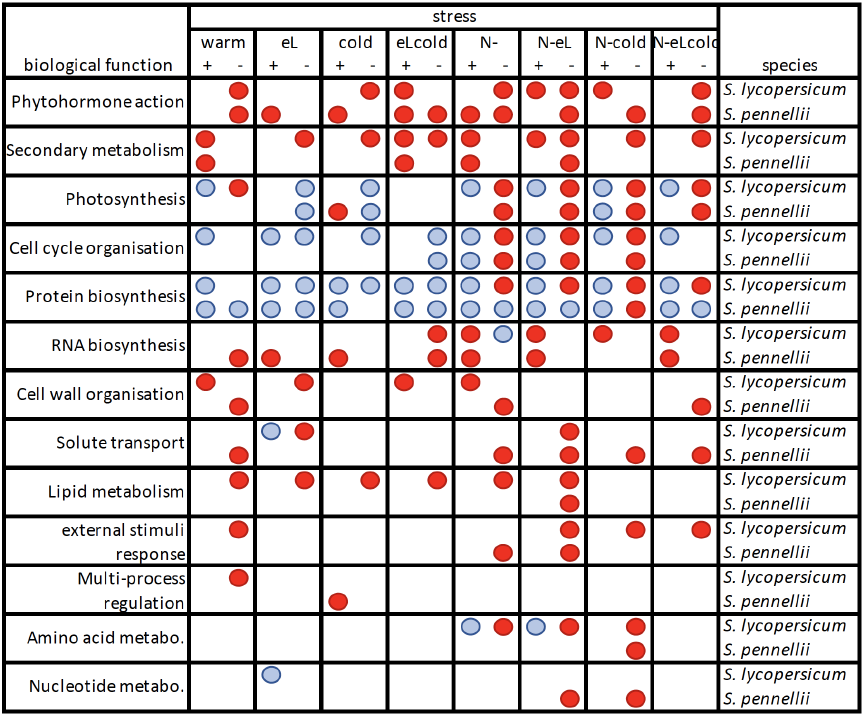
Results of an overrepresentation analysis (ORA) performed for *S. lycopersicum* and *S. pennellii*. Plants were grown under different abiotic stress conditions (warmer or cooler temperature regime: warm or cold, elevated light intensities: eL, nitrogen deficiency: N-, and combinations thereof as indicated) on rockwool. The over- or underrepresentation (highlighted in red and blue dots, respectively) of indicated Mercator bins (Lohse et al., 2014; Usadel et al., 2009; Schwacke et al., 2019) were calculated seperately for up- (+) and down-regulated genes (-) using a Fisher test (Usadel et al., 2006) in combination with a multiple testing correction as performed by Benjamini and Hochberg (1995).

In this respect, pathways belonging to phytohormone action and secondary metabolism were induced in case of upregulated genes, and repressed in case of downregulated genes under various stress treatments in both species. Induction and repression can be observed at the same time (e.g. phytohormone action in *S. lycopersicum* for N-eL), which is not a contradiction, as the shown main categories in table 2 comprise different sub-pathways. Therefore, induction of one pathway and repression of another may result in an overrepresentation of the main category in both up- and down-regulated genes.

Interestingly, the category of protein biosynthesis was repressed under any condition harboring nitrogen deficiency for *S. lycopersicum*, while *S. pennellii* was not responding, with the exception of N-cold conditions. On the other hand, photosynthesis pathways and cell cycle organisation were indicated as being repressed under any kind of nitrogen deficiency treatment for both species. RNA biosynthesis pathways were often induced in both species, but not always under the same abiotic conditions. Pathways of other categories, like cell wall organisation, solute transport, lipid metabolism, external stimuli, multi-process regulation, or metabolism of amino acids or nucleotide, were more often repressed in the two species.

Besides the general categories in the ORA, we also investigated specific category bins (see supplementary figure 4 and 5) with respect to the applied abiotic stress conditions.

Under **elevated light intensities**, the cultivated tomato *S. lycopersicum* responded with induction of genes involved in photosynthesis system II and chlororespiration in combinaion with repressed genes involved in biosynthesis of lignin and hydroxyproline-rich glycoproteins (see supplementary figure 4). In addition, the cultivated tomato repressed genes involved in S-glutathionylation of proteins, lipid degradation, secondary metabolism (biosynthesis of aurones), RNA biosynthesis regulated via myeloblastosis (MYB)- or Asymetric Leaves2/Lateral Organ Boundaries (AS2/LOB)-transcription factors, and solute transport via the ATP-binding cassette (ABC) transporter superfamily.

For *S. pennellii* genes involved in abscisic acid (ABA), cytokinin, and jasmonic acid perception and signalling, as well as other signalling peptides were induced, together with pathways belonging to RNA biosynthesis regulated via MYB-, Apetala2 / Ethylene Response Factor (AP2/ERF)-, NAC (NAM, ATAF and CUC)-, WRKY- or TIFY-transcription factors (see supplementary figure 5). In addition, genes involved in preinitiation of DNA replication, RNA biosynthesis regulated via Plant Homeo Domain (PHD)-transcription factors, or protein phosphorylaion via PEK were repressed in the wild tomato.

Under **warmer temperature**, *S. lycopersicum* induced pathways of terpene and aurone biosynthesis, transcriptional regulation of RNA biosynthesis by MYB- and AP2/ERF-transcription factors, solute transport via the ATP binding cassette (ABC) transporter superfamily, as well as pathways belonging to cell wall organisation by hydroxyproline-rich glycoproteins and cuticular lipid formation (see supplementary figure 4), besides small heat shock proteins (HSP). Pathways of the photosynthesis (photosystem II phosphorylation, ATP synthesis, calvin cycle), tetrapyrrol biosynthesis, preinitiation of DNA replication, ribosomal protein biosynthesis and protein S-glutathionylation were repressed. In the main category cellular respiration, pathways belonging to oligosaccharide metabolism and amino acid degradation were induced, while pathways of desaturation of fatty acids were repressed. In addition in the main category phytohormone action, genes for auxin conjugation and degradation were induced, while genes belonging to gibberellin signaling were repressed.

Under these conditions the wild relative *S. pennellii* induced pathways belonging to the biosynthesis of p-coumaroyl-CoA (a precursor of phenolics) and phytosterol (components of the membrane lipid bilayer), and HSP (see supplementary figure 5). Protein modification via tyrosine kinase-like (TKL) superfamily was induces, whereas pathways of S-glutathionylation were repressed. In addition, transcriptional regulation of RNA biosynthesis was repressed under warmer temperatures, as well as cell wall organisation in the context of cellulose-hemicellulose network assembly, lignol conjugation and polymerisation, and pectin modification. In the main category phytohormone action, pathways of brassinosteroid conjugation and degradation were induced in addition to signalling pathways via non-cystein rich peptides (NCRP), while signalling pathways via cystein rich peptides (CRP) were repressed. Carrier-mediated solute transport and solute transport via major intrinsic protein (MIP) family channels were also repressed.

Under **chilling temperature** conditions only minor changes were observed for *S. lycopersicum*. Here, pathways of pectin modification and degradation were induced (see supplementary figure 4). Whereas, lipid desaturation and pathways belonging to preinitiation of DNA replication, protein biosynhthesis and S-gluthathionylation, tetrapyrrol biosynthesis, and gibberellin signaling were repressed, besides pathways for the solute transport via drug/metabolite transporter (DMT) superfamily transporter.

For *S. pennellii* more changes were detected (see supplementary figure 5). Pathways belonging to photosynthesis (phosphorylation, photosystem II, chlororespiration), RNA biosynthesis mediated by MYB-, AP2 / ERF-, and TIFY-transcription factors were induced. In contrast, RNA biosynthesis mediated by PHD-transcription factors and pathways belonging to RNA processing (3’ end processing, splicing, surveillance), microfillament organisation, vesicle trafficking, protein translocation, and solute primary active transport via P-type ATPase superfamily were repressed. Pathways of ribosomal protein biosynthesis were repressed, whereas mitochondrial and platstidial protein biosynthesis were induced. Phosphorylation of proteins by atypical protein kinase families were repressed, while S-gluthathionylation of proteins was induced. Protein quality control by small HSP were induced, while ubiquitin-conjugation was repressed. In the main category lipid metabolism, pathways of lipid-bodies associated activities were induced, whereas pathways of lipid trafficking were repressed. Gibberellin- and auxin-mediated pathways in the main category phytohormone action were repressed, while signaling mediated by NCRP and CRP were induced in this main category. We observed an induction of histone pahways, whereas genes involved in histone deacetylation and chromatin remodeling complex were repressed.

Under a **combination of chilling temperatures and elevated light intensities**, we observed, that induced pathways belonging to photosynthesis under the single stress of elevated light intensities in *S. lycopersicum*, were now repressed (see supplementary figure 4). In addition, pathways belonging to lipid degradation, transcriptional regulation of RNA biosynthesis via MYB-, homeobox-, AP2/ERF-, NAC-, WRKY-, and TIFY-transcription factors, cell wall organisation via hydroxyproline-rich glycoproteins, solute transport via ABC superfamily transporter were repressed. Also, genes involved in mitochondrial and plastidial protein biosynthesis, as well as phosphorylation of proteins via TKL protein superfamily, S-glutathionylation of proteins and glutaredoxins were repressed. In the main category of secondary metabolism, an ambivalent pattern for pathways of flavonoid biosynthesis was observed with being partly repressed and partly induced, while aurone biosynthesis was repressed. Auxin, cytokinin and jasmonic acid conjugation and degradation were repressed, while phytohormone signaling via NCRP was activated.

In *S. pennellii* genes involved in phytosterol biosynthesis, terpene biosynthesis, and xylan modification were induced (see supplementary figure 5). Whereas, genes involved in RNA silencing, protein phosphorylation via Ca^2+^/calmodulindependent protein kinase (CAMK) superfamily, glutaredoxins, response to blue light, and solute transport via MIP family channels were repressed. In the main category phytohormone actions, cytokinin conjugation and degradation, as well as jasmonic acid conjugation and degradation, and NCRP were induced, while absisic acid perception and signalling, and ethylene biosynthesis were repressed. Regulation of RNA biosynthesis via C2C2- and MYB-transcription factors was down-regulated, while regulation via AP2/ERF-transcription factors was up-regulated.

**Nitrogen deficiency** showed the strongest effects in both species, and the induction / repression of genes was often consistent in samples of stress treatments harboring nitrogen deficiency (see supplementary figure 4 and 5). The following description reflects all nitrogen deficiency harboring treatments if nothing else is mentioned.

Analysis of subcategories revealed a strong repression of photosynthesis pathways in *S. lycopersicum* (see supplementary figure 4), accompanied by a repression of deoxynucleotide biosynthesis, amino acid biosynthesis, tetrapyrrol biosynthesis, redox homeostasis (only for nitrogen deficiency as single stress and the triple stress N-eLcold), solute transport via DMT superfamily and MIP family channels. In addition chromatin organisation via histones was repressed, as well as DNA replication pathways, protein biosynthesis, HSP biosynthesis (except N-eL) and glutaredoxins. In contrast pathways of cuticular lipid formation and protein phosphorylation via TKL protein kinase superfamily (only N- and N-eL) were induced. Overall, regulation of RNA biosynthesis via MYB-, homeobox- (only nitrogen deficiency as single stress), GRAS- (Gibberellic-acid insensitive (GAI), Repressor of GAI (RGA), and Scarecrow (SCR)), and GRF-GIF (Growth-Regulating Factor and GRF-Interacting Factor)-transcription factors were induced, while signaling via AP2/ERF transcription factor was repressed (only in triple stress N-eLcold). In the main category cellular respiration, pathways belonging to pyruvate production were induced (except for nitrogen deficiency as single stress), while pathways of aspartate biosynthesis and phospholipase activities were repressed (except for the triple stress N-eLcold). In addition, for the pathways of the secondary metabolism we observed an induction of flavonoid biosynthesis and a repression of terpene synthase. Also for phytohormone action, we observed a diverse pattern, with pathways for auxin eflux (only in N-eLcold treatment), NCRP protein signaling (only N-eL), and gibberelin signaling being induced, while pathways of cytokinin conjugation and degradation (only N-eL and triple stress N-eLcold), NCRP signaling (except N-cold) and CRP signaling were repressed.

In the wild relative *S. pennellii*, repression of photosynthesis was accompanied by repression of pathways for ammonium assimilation (except for N-eLcold), and pathways of amino acid, deoxynucleotide, and tetrapyrrol biosynthesis (see supplementary figure 5). In addition, chromatin organisation via histones (except triple stress N-eLcold), DNA replication, cytoskeleton organisation (N- and N-eL), cell wall organisation via pectin modification were repressed. In contrast, transcriptional regulation of RNA biosynthesis via MYB-, homeobox- (only N- and N-eL), AP2/ERF- (except the triple stress), WRKY-, and GRF-GIF-transcription factor were induced, as well as pathways belonging to protein modification via TKL superfamily (only N-cold), S-glutathionylation and glutaredoxins. Pathways belonging to protein biosynthesis were repressed under N-cold, while protein translocation to chloroplasts was repressed under nitrogen deficiency. Pathways for HSP biosynthesis were induced under N-eL, while proteolyse via serine-type peptidase (N- and N-eL) and aspartic-type peptidase (N- and N-eLcold) were repressed. For subcategories of the lipid metabolism we found a more diverse pattern with repressed pathways of phosphatidylcholine (only N-eL) and induced pathways for lipid bodies-associated activities. In the categories of secondary metabolism, pathways belonging to terpenoid biosynthesis were repressed (only N-cold), while pathways belonging to flavonoid biosynthesis were induced (except for N-cold). In the main category of phytohormone action absisic acid signaling (except for N-single stress), cytokinin signaling (only N-eL) and NCRP signaling (only N- and N-cold) were repressed, while NCRP signaling was induced under N-eL and the triple stress N-eLcold. Sub-categories of solute transport were mainly repressed (carrier-mediated transport as well as MIP family channels).

### 3.3 Co-Expression networks for nitrogen deficiency and chilling temperatures

With the help of a weighted cluster analysis (Langfelder and Horvath, 2008) we identified co-expression networks (modules) related to a specific stress (binary trait). The network matrix was built by checking for differences between the control group and the stress treatments. In addition, single stresses were set as binary trait.

For each module the correlation for all harboring genes with the binary trait (warm, cold, elevated light intensities, or nitrogen deficiency) was calculated. In total we identified 41 modules for *S. lycopersicum* (see supplementary figure 6) and 44 modules for *S. pennellii* (see supplementary figure 7). Modules with a correlation of 0.72 and higher (later called significant modules) for a specific binary trait were investigated further to identify characteristics among those genes.

First, for all genes within a significant module we checked the presence of up- and down-regulated genes for each applied stress condition (see supplementary figures 8 and 9). Significant modules for a binary trait are enriched for genes that respond to the binary trait by induction or repression. Therefore, significant modules show a higher percentage of up- or down-regulated genes under a stress condition harboring the binary trait compared to stress conditions without the binary trait. In addition, significant modules are enriched for genes stronger induced or repressed under a stress condition harboring the binary trait, as indicated by higher absolute maximal and minimal log fold change (FC). Data from the whole genome was used as a control group for comparison.

For *S. lycopersicum* we investigated the modules brown, midnightblue and white (all related with warm), grey60 (related with cold), and purple (related with nitrogen deficiency) (see supplementary figure 6). They included between 130 and 16877 genes. For all modules but module white, we found an enrichment for up-regulated genes under the respective stress (see supplementary figure 8). In addition, the absolute value of maximal / minimal log FC increased respectively, for the correlating stresses when compared with other stress conditions, indicating that the changed expression responses will have a higher biological relevance and impact on the stress response.

We observed a similar pattern for the modules darkred, blue, darkturquoise (all related with nitrogen deficiency) and purple (related with cold) for *S. pennellii* (see supplementary figure 9), although here some deviation was found between the treatments (e.g. N-cold showed less up-regulated genes in module darkred and blue). Those modules harbored between 449 and 6278 genes. Still, we conclude, that accumulated genes within the modules respond mainly to the respective trait.

#### Nitrogen deficiency induces secondary metabolism

For both species, we focused on genes of the modules that are associated with nitrogen deficiency (module purple for *S. lycopersicum* and module blue for *S. pennellii*). The 40 up-regulated genes of highest significance (p<0.01) for each stress condition harboring nitrogen deficiency were checked for their (putative) biological function. In total almost half (21 genes) of the top 40 genes were up-regulated under all stress conditions harboring nitrogen deficiency for *S. lycopersicum* and 42.5 % (17 genes) in *S. pennellii* (see figure 2).

**Fig. 2:**
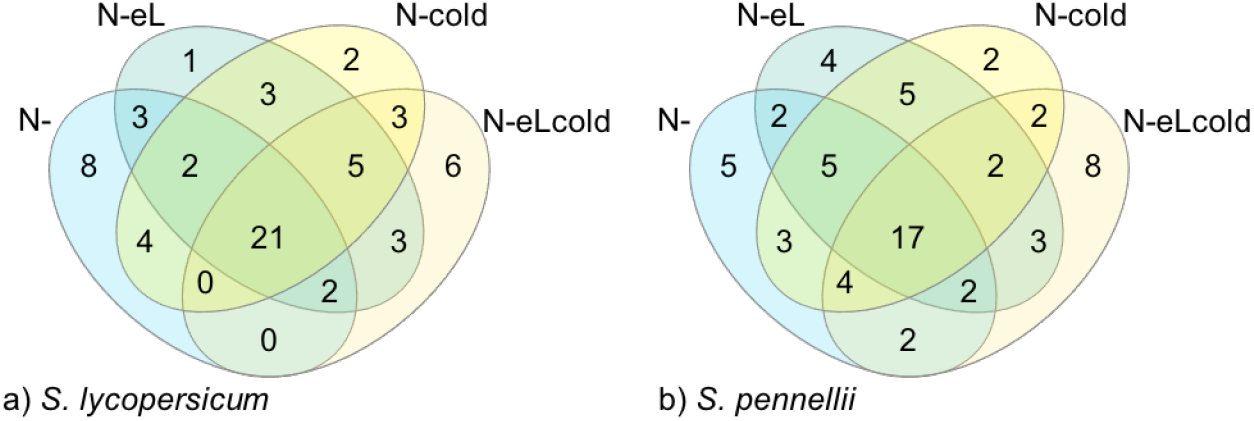
Venn diagram showing the overlap for top 40 up-regulated genes in modules related with nitrogen deficiency. Up-regulated genes in the module purple (*S. lycopersicum*, a) and blue (*S. pennellii*, b) were analysed for their expression pattern under stresses harboring nitrogen deficiency. The top 40 genes were intersected to identify common genes involved in the stress response under all conditions harboring nitrogen deficiency.

For *S. lycopersicum* we found in total 55 different genes among the top 40 genes, that were up-regulated in at least one nitrogen deficiency harboring stress condition (see supplementary table 1). 5 genes of unknown function (out of 15) were up-regulated under all four stress conditions (see table 3), as was a putative remorin (Solyc05g014710.4.1) and a chalcone synthase (Solyc05g053550.3.1). The last indicates a possible induction of phenolics biosynthesis. Another gene of this pathway, the isoflavone reductase homolog (Solyc10g052500.2.1) is up-regulated only under N-cold condition. Interestingly, three additional not further characterised putative glycosyltransferases could be determined (see table 3).

**Table 3:** Common (putative) genes of the modules purple (*S. lycopersicum*) and blue (*S. pennellii*) among the top 40 highly significant up-regulated genes in all stresses harboring nitrogen deficiency. For each indicated stress the top 40 genes of the module purple (*S. lycopersicum*) and module blue (*S. pennellii*) were checked for their biological (putative) function.

| species | gene ID | (putative) function | N- | N-eL | N-cold | N-eLcold |
| --- | --- | --- | --- | --- | --- | --- |
| *S. lycopersicum* | Solyc05g053550.3.1 | chalcone synthase | + | + | + | + |
|  | Solyc10g052500.2.1 | Isoflavone reductase homolog |  |  | + |  |
|  | Solyc05g014710.4.1 | putative Remorin | + | + | + | + |
|  | Solyc04g082860.3.1 | putative UDP-glycosyltransferase |  | + |  | + |
|  | Solyc10g085880.1.1 | putative UDP-glycosyltransferase |  | + |  | + |
|  | Solyc04g074340.3.1 | putative UDP-glycosyltransferase |  |  | + |  |
|  | Solyc01g098420.3.1 | unknown function | + | + | + | + |
|  | Solyc02g031730.1.1 | unknown function | + | + | + | + |
|  | Solyc04g016085.1.1 | unknown function | + | + | + | + |
|  | Solyc06g068130.3.1 | unknown function | + | + | + | + |
|  | Solyc07g008210.4.1 | unknown function | + | + | + | + |
| *S. pennellii* | Sopen10g032910.1 | flavonol-3-O-rhamnosyltransferase |  |  |  | + |
|  | Sopen03g025580.1 | Mlo family | + | + | + | + |
|  | Sopen11g003340.1 | putative UDP-glycosyltransferase | + | + | + | + |
|  | Sopen03g021390.1 | putative UDP-glycosyltransferase | + |  |  |  |
|  | Sopen06g025400.1 | unknown function | + | + | + | + |
|  | Sopen07g004090.1 | unknown function | + | + | + | + |
|  | Sopen08g006790.1 | unknown function | + | + | + | + |
|  | Sopen11g007320.1 | unknown function | + | + | + | + |
|  | Sopen11g017460.1 | unknown function | + | + | + | + |

For *S. pennellii* we found 66 different genes among the top 40 genes, that were up-regulated in at least one nitrogen deficiency harboring stress condition (see supplementary table 1). 19,6 % (17 genes) were of unknown function, and out of those five were up-regulated under all four conditions (see table 3). 47,0 % (31 genes) were of putative function, and out of these Sopen03g025580.1 (putative Mlo family) and Sopen11g003340.1 (putative UDP-glycosyltransferase) were up-regulated under all conditions. Interestingly, we found an additional gene with a putative UDP-glycosyltransferase function (Sopen03g021390.1) up-regulated under nitrogen deficiency. The only gene belonging to the secondary metabolism Sopen10g032910.1 (flavonol-3-O-rhamnosyltransferase) was found in the top 40 genes under the triple stress of nitrogen deficiency, elevated light intensities and chilling temperatures (see table 3).

We performed a BLAST analysis with the putative UDP-glycosyltransferase (UGT) genes and the unknown genes as indicated in table 3 of both species to get some further insights on their function (for sequences see supplementary table 1). The multiple alignment revealed some homology, including the common “Plant secondary product motif” (PSPG) for the **putative UGT** genes (Gachon et al., 2005) within each species and also between them. In addition, we searched for homologous genes with more than 50 % query coverage and 75 – 100 % sequence identity.

For the two genes with putative UGT function from *S. pennellii* all hits belong to the family of Solanaceae. Most of the hits are of unknown function, but for Sopen11g017460.1 three hits are described to encode for an UGT in *Nicotiana attenuata* (99 % identity for each hit, see also supplementary file 1). Most of the found hits for *S. lycopersicum* encode for genes with yet undefined function (see also supplementary table 1). But for Solyc04g074340.3.1 the hit from *Lycium barbarum* encodes for an UGT gene (85 % identity). In addition, for Solyc10g085880.1.1 three hits from *Nicotiana attenuata* (77-82 % identity) encode as well for an UGT gene.

We performed the same kind of analysis for genes with yet **unknown function** (multiple alignment and single sequence BLAST analysis with 50 – 100 % query coverage and 75 – 100 % identity). These genes shared some minor sequence homology but only pair-wise.

For *S. lycopersicum*, our analysis of single BLAST results for each of the afore-mentioned genes gave only hits that belong to the family of Solanaceae. Besides hits from *S. lycopersicum* and *S. pennellii*, we found mainly hits belonging to cultivated or wild potato (*Solanum tuberosum* or *Solanum pinnatisectum*). The same was true for hits belonging to genes from *S. pennellii*. But for Sopen11g007320.1, we found additional hits belonging to Fagaceae, Malvaceae, Rosaceae as well as *Arabidopsis thaliana*. For all genes of unknown function, none of the found hits were further characterised for a biological function.

#### Response to chilling temperatures is more diverse

In both species we identified modules related with the binary trait of chilling temperatures. In case of *S. lycopersicum* we investigated the module grey60 (n = 982) and for *S. pennellii* module purple (n = 736) to identify genes related with the stress response to chilling temperatures. The genes within the modules were analysed regarding their expression pattern (p<0.02) under different stress combinations harboring chilling temperatures.

When screening the expression pattern for *S. lycopersicum* we noticed that genes were either up-regulated under chilling temperatures or a combination of elevated light intensities and chilling temperatures, or down regulated in combined stresses harboring nitrogen deficiency and cold (N-cold and N-eLcold) (see supplementary table 1, and figure 3 a). Therefore, we focused on the common genes under chilling temperatures or a combination of elevated light intensities and chilling temperatures (see table 4). Here we found seven genes (including two genes with unknown function) belonging to cell cycle organisation (Solyc02g079370.3.1), lipid metabolism (Solyc08g063080.3.1), and phosphorylation of proteins (Solyc12g009190.2.1), as well as a cytochrome P450 monoxygenase (Solyc03g111950.3.1) and a calcium sensor protein (Solyc12g055920.3.1).

**Fig. 3:**
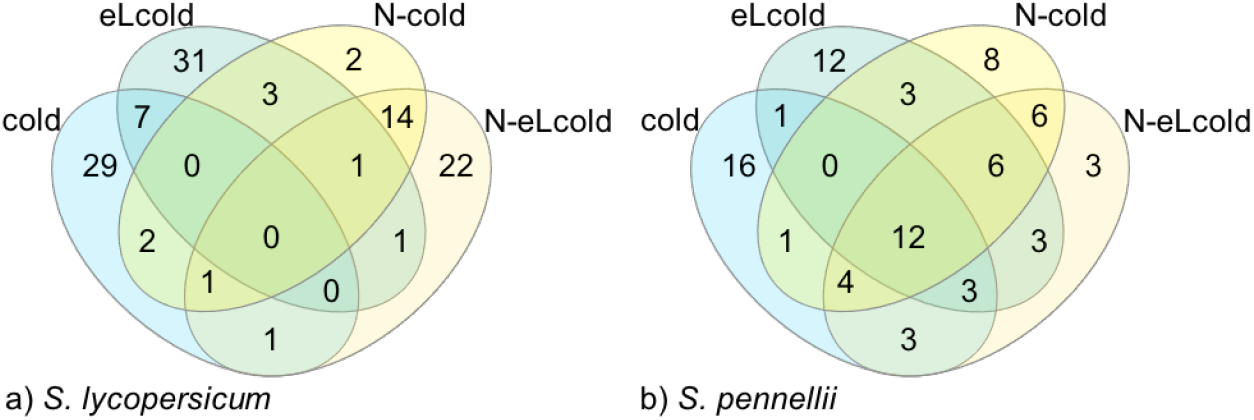
Venn diagram showing the overlap for top 40 genes in modules related with chilling temperatures. Genes in the module grey 60 (*S. lycopersicum*, a) or purple (*S. pennellii*, b) were analysed for their expression pattern under stresses harboring chilling temperatures. The top 40 genes were intersect to identify common genes involved in the stress response under all conditions.

**Table 4:** Common genes under chilling temperatures. Top 40 up-regulated genes were investigated concerning their expression pattern and biological function under stress conditions harboring chilling temperatures.

| species | gene ID | (putative) function | cold | eLcold | N-cold | N-eLcold |
| --- | --- | --- | --- | --- | --- | --- |
| *S. lycopersicum* | Solyc02g079370.3.1 | cyclin D 6;3 | + | + |  |  |
|  | Solyc03g111950.3.1 | Cytochrome P450 | + | + |  |  |
|  | Solyc08g063080.3.1 | putativ UDP-sulfo-quinovose synthase | + | + |  |  |
|  | Solyc12g009190.2.1 | protein kinase (LRR-III) | + | + |  |  |
|  | Solyc12g055920.3.1 | Calcineurin B-like protein 4 | + | + |  |  |
|  | Solyc02g067480.3.1 | unknown function | + | + |  |  |
|  | Solyc09g008270.3.1 | unknown function | + | + |  |  |
| *S. pennellii* | Sopen03g022310.1 | HD-ZIP I/II transcription factor | + | + | + | + |
|  | Sopen04g006840.1 | Fumarylacetoacetate (FAA) hydrolase family | + | + | + | + |
|  | Sopen08g003230.1 | HD-ZIP I/II transcription factor | + | + | + | + |
|  | Sopen09g035000.1 | sucrose-phosphate synthase | + | + | + | + |
|  | Sopen01g044210.1 | unknown function | + | + | + | + |
|  | Sopen02g015400.1 | unknown function | + | + | + | + |
|  | Sopen02g019230.1 | unknown function | + | + | + | + |
|  | Sopen02g026300.1 | unknown function | + | + | + | + |
|  | Sopen03g038840.1 | unknown function | + | + | + | + |
|  | Sopen05g029120.1 | unknown function | + | + | + | + |
|  | Sopen06g017630.1 | unknown function | + | + | + | + |
|  | Sopen06g033440.1 | unknown function | + | + | + | + |

For *S. pennellii* we found a more homogenous expression pattern for genes clustered in module purple, so that many genes were up-regulated in any combination of stresses harboring chilling temperatures (see supplementary table 1, and figure 3 b). When intersecting the top 40 significantly up-regulated genes, we identified 12 common genes present in all stress conditions (see table 4), among which eight are of yet unknown function.

We performed a sequence analysis for genes with unknown function (see table 4) of both species. While performing a multiple alignment with all genes, no similarity between or within the species could be identified. Most hits of a BLAST analysis (50 – 100 % query coverage, 75 – 100 % identity) belong to the Solanaceae family (besides *S. lycopersicum* and *S. pennellii*, also potato *Solanum tuberosum* and *Solanum pinnatisectum*, tobacco *Nicotiana benthamiana*, or boxthorn *Lycium chinense*). And almost none of them were further characterised for a specific function. Only for Solyc09g008270.3.1, hits belonging to *Arabidopsis thaliana* (At2g36885) were found, that encodes for a translation initiation factor. In addition for Sopen02g015400.1, hits belonging to *Nicotiana benthamiana* or *Lycium chinense* encode for a salicylic acid binding protein.

In addition, we were able to identify three related modules in the clustered network analysis for *S. lycopersicum* associated with the binary trait of warmer temperatures (modules brown, midnightblue and white, see supplementary figure 6). Due to higher significance and enrichment of up-regulated genes, we investigated only the module brown further (see also supplementary table 1).

Module brown comprised 2098 genes. Among the 40 up-regulated genes, 2 HSP genes were identified (Solyc01g102960.3.1 and Solyc11g020330.1.1), three genes belong to cell wall organisation (Solyc06g060970.2.1, Solyc08g077900.3.1 and linebreak Solyc12g088240.2.1), two genes to amino acid metabolism (Solyc08g076980.4.1 and Solyc06g007180.3.1), and three further genes to solute transport (Solyc03g019820.3.1, Solyc06g072130.4.1 and Solyc06g051860.3.1). Six genes were of unknown function, and 18 genes were only of putative function.

### 3.4 The tomato species differ in their stress induced accumulation of metabolites

A couple of studies investigated the partly volatile metabolite composition of either tomato fruits (Tieman et al., 2006, 2012) or trichomes (Schilmiller et al., 2012; Leong et al., 2019), and in addition a comparison of the metabolite composition in leaf samples from a cultivated and a wild tomato species were reported for adult plants (Schauer et al., 2004; Tohge et al., 2020), but has not been conducted so far by applying mild abiotic stress conditions.

Therefore, we complemented our analysis with a metabolite analysis using a combined LC-FTICR-MS and LC-MS/MS approach. In a first step highly accurate masses were obtained by LC-FTICR-MS allowing the calculation of chemical formulae. In a second step these precursor ions were submitted to LC-MS/MS in order to get their daughter ion spectra for structure elucidation. For *S. lycopersicum* we could identify six enriched secondary metabolites using the ESI(+) mode when compared to the control (see figure 4 and supplementary table 2), and additional six primary metabolites when detection was performed in the ESI(-) mode (see figure 5 and supplementary table 2).

**Fig. 4:**
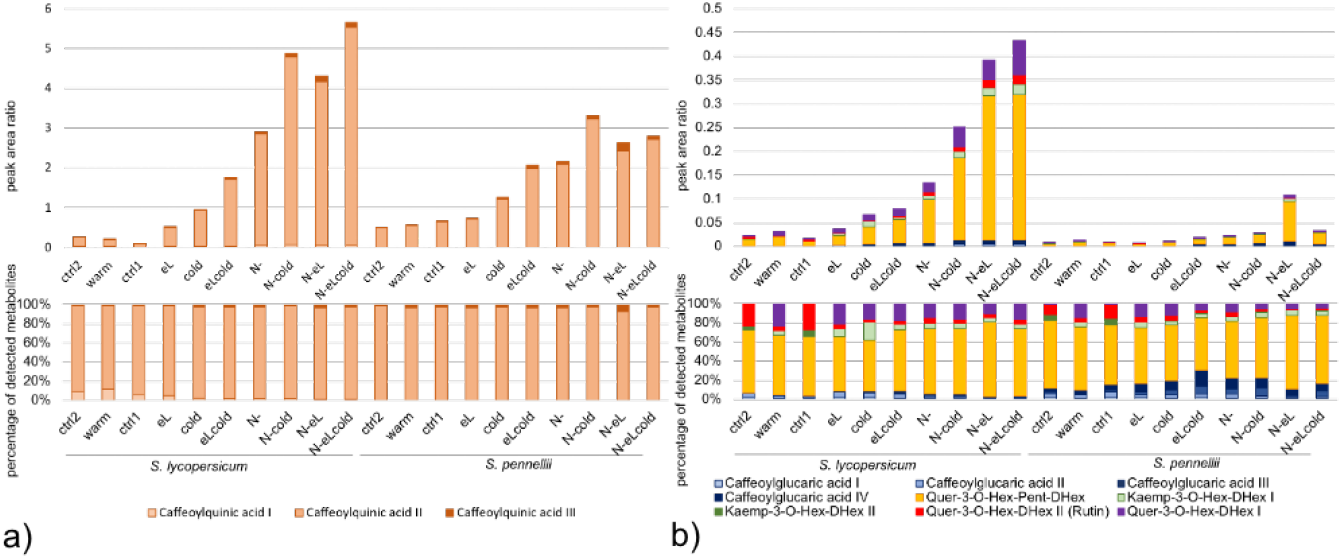
The cultivated tomato accumulated more secondary metabolites than the wild species (obtained by ESI(+) mode). Samples used for the RNASeq analysis and the qPCR verification were used for determination of secondary metabolites by LC-ESI(+)-FTICR-MS and LC-ESI(+)-MS/MS. **(a)** Monocaffeoylquinic acid (CQA) (3 regioisomers, I - III), and **(b)** caffeoylglucaric acid (4 regioisomers, I - IV), Quer-3-O-Hex-Pent-DHex, Kaemp-3-O-Hex-DHex (2 regioisomers, I and II), rutin and Quer-3-O-Hex-DHex I were unambiguously identified. Quantification of them was performed with LC-MS/MS by use of their corresponding MRM transitions. Peak area ratios (above) and their normalized illustration (below) are shown as mean of two independent biological experiments.

**Fig. 5:**
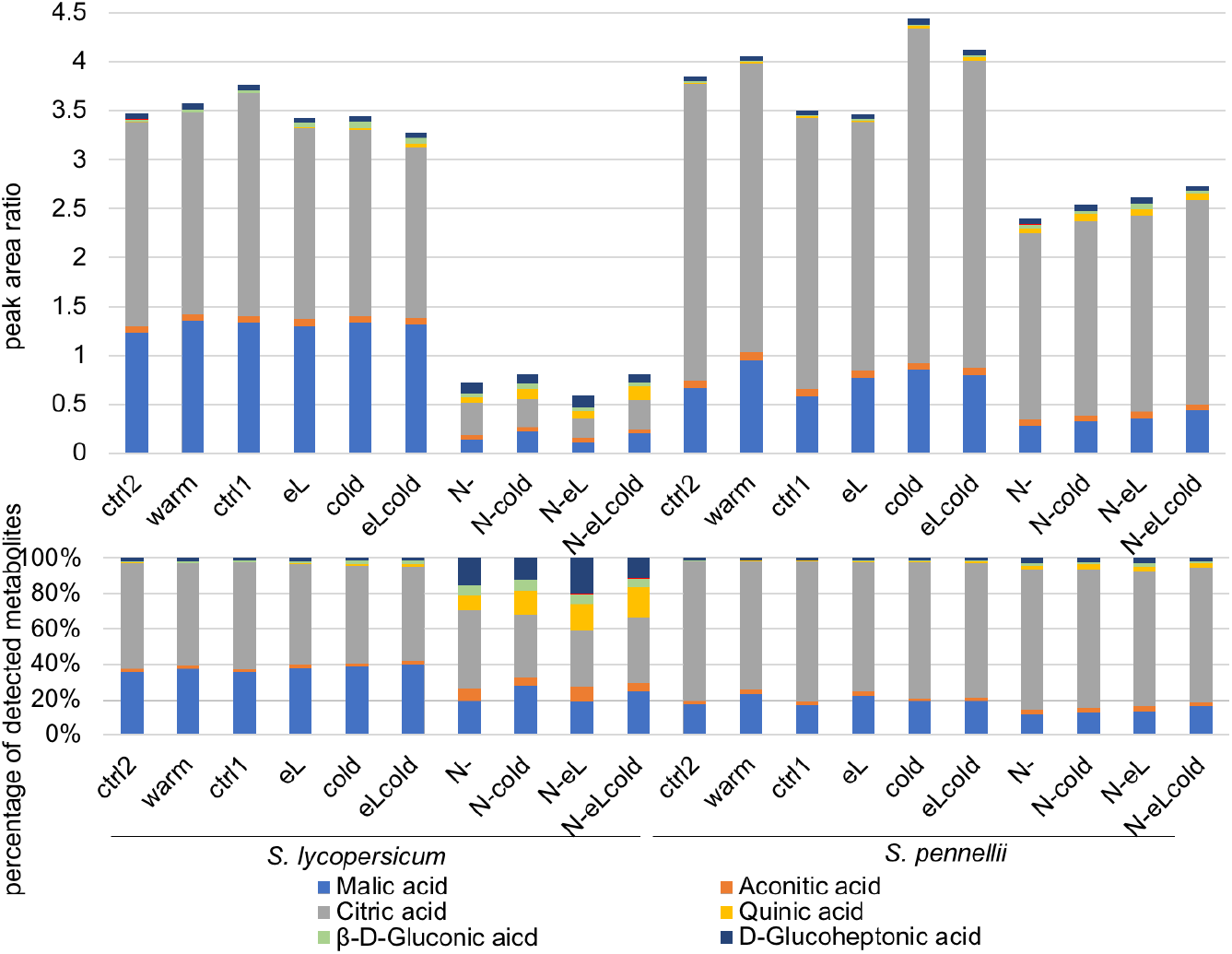
The two tomato species responded similar for a set of primary metabolites obtained by ESI(-) mode. Samples used for the RNASeq analysis and the qPCR verification were also used for determination of metabolites by LC-ESI(-)-FTICR-MS and LC-ESI(-)-MS/MS. Malic acid, aconitic acid, citric acid, quinic acid, gluconic acid and glucoheptonic acid were unambiguously identified. Quantification of them was performed with LC-MS/MS by use of their corresponding MRM transitions. Peak area ratios (above) and their normalized illustration (below) are shown as mean of two independent biological experiments.

Mono-caffeoylquinic acid (CQA) was found with highest amounts in both species (see figure 4 a and supplementary table 2). Caffeoylquinic acid is a group of compounds composed of a quinic acid core, acylated with one or more caffeoyl groups. Among the best described members of this group is chlorogenic acid (5-O-caffeoylquinic acid), a mono-caffeoylquinic acid (Liu et al., 2020). We detected three different regioisomers of CQA, while only one of them was produced in high amounts (more than 93 %, see supplementary table 2). With our approach we were unable to further elucidate the structural nature of the detected regioisomers, but it is highly probable that QCA II is equivalent to chlorogenic acid.

Both species show only a low induction under warmer temperature and elevated light intensities (see also supplementary figure 10). For chilling temperatures as well as a combination of elevated light intensities and chilling temperatures the amount of CQA rose. The highest quantities were found for nitrogen deficiency and any combinatorial stress harboring nitrogen deficiency.

In addition, in lower amounts five other metabolites were detected (see figure 4 b), namely caffeoylglucaric acid (4 regioisoforms), quercetin-3-O-deoxyhexosyl-pentosyl-hexoside (Quer-3-O-Hex-Pent-DHex), kaempferol-3-O-deoxyhexosyl-hexoside (Kaemp-3-O-Hex-DHex, two isoforms), quercetin-3-O-deoxyhexosyl-hexoside II (rutin, Quer-3-O-Hex-DHex II), and quercetin-3-O-deoxyhexosyl-hexoside (Quer-3-O-Hex-DHex). Overall the induction in *S. lycopersicum* for those five metabolites followed the induction of CQA, showing the highest response in any combinatorial stress harboring nitrogen deficiency. Among these Quer-3-O-Hex-Pent-DHex and Quer-3-O-Hex-DHex II (rutin) were substantially induced in *S. lycopersicum* (see figure 4 b).

In contrast, *S. pennellii* did not show such a strong response of those five metabolites for any stress with the exception of the combination of nitrogen deficiency and elevated light intensities. In addition, the ratio inbetween the five metabolites was changed, when compared with *S. lycopersicum*. Here, more caffeoylglucaric acid was produced than Quer-3-O-Hex-Pent-DHex.

The regioisomers Kaemp-3-O-Hex-DHex I and Quer-3-O-Hex-DHex I showed the highest induction rate (see also supplementary figure 10 a) in both species, but the absolute abundance of both metabolites is low within the samples. While the induction rate of the regioisomers Kaemp-3-O-Hex-DHex II and Quer-3-O-Hex-DHex II (rutin) were the mildest of all analysed metabolites obtained by ESI(+) mode. In total, the induction rate of nitrogen deficiency as single stress was always higher than for other single stresses (eL, warm or cold) in *S. lycopersicum*, and the induction rate of combined stress conditions were often higher, than for the single stress conditions in both species (see also supplementary figure 10 a).

*S. lycopersicum* induced the metabolite production much stronger than *S. pennellii*. As an example, for CQA II the induction rate was 59.4-fold in *S. lycopersicum* (for N-eLcold), and 4.9-fold in *S. pennellii* (for N-cold), and for Quer-3-O-Hex-Pent-DHex we observed a 30.6-fold induction in *S. lycopersicum* (N-eL and N-eLcold) and a 14.6-fold induction in *S. pennellii* (N-eL) (see also supplementary figure 10 a).

By using the ESI(-) mode we investigated six organic acids mainly coming from the primary metabolism (see figure 5), namely malic acid, aconitic acid, citric acid, quinic acid, *β*-D-gluronic acid, and D-glucoheptonic acid.

A strong reduction of malic acid and citric acid were observed for all stresses harboring nitrogen deficiency. This was more prominent in *S. lycopersicum* than in the wild species *S. pennellii* (see also supplementary figure 10 b), indicating that the primary metabolism will be restricted under nitrogen deficiency. In addition, quinic acid, *β*-D-gluronic acid, and D-glucoheptonic acid were induced up to 44-fold in *S. lycopersicum* (quinic acid, N-eLcold) and up to 6-fold in *S. pennellii* (quinic acid, N-eLcold) (see supplementary figure 10 b). As a result, the ratio between the metabolites changed under these condition, so that citric acid and malic acid are less prominent, and quinic acid, *β*-D-gluronic acid, and D-glucoheptonic acid increase relatively in both species (see figure 5 bottom).

Overall, nitrogen deficiency in any stress combination resulted in a strong induction of secondary metabolites for the cultivated tomato and a reduction of primary metabolites in both species. Thereby, the pattern of a stronger response under nitrogen deficiency is complementing our RNASeq data.

### 3.5 Structural pathways for secondary metabolism biosynthesis in cultivated and wild tomato

To gain further insights in the time-dependent induction and to validate the metabolite composition we performed quantitative real time expression analysis with samples harvested four and seven days after induction of stress treatments. We chose structural genes involved in the flavonoid biosynthesis pathway (see also figure 7) belonging to the bins for flavonoid biosynthesis in the overrepresentation analysis. In addition we chose for two structural genes of the anthocyanidin biosynthesis belonging to the bins for anthocyanidin biosynthesis in the overrepresentation analysis. All chosen genes were also members of the nitrogen deficiency related modules purple (*S. lycopersicum*) and blue (*S. pennellii*) (see supplementary table 1). With that approach RNASeq results data could be verified (see figure 6).

**Fig. 6:**
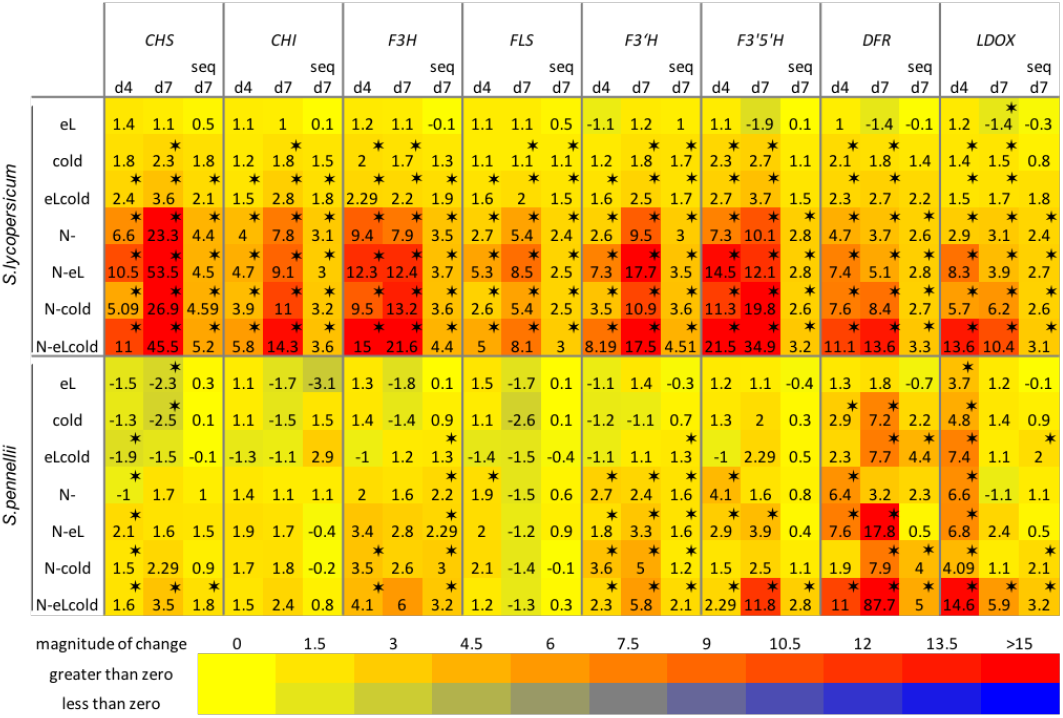
Expression of structural genes involved in secondary metabolism. RNASeq data were verified using samples from an independent biological experiment performed within the same plant chambers under the same stress conditions as before. *S. lycopersicum* and *S. pennellii* were grown on rockwool for six weeks before applying chilling temperatures (cold), elevated light intensities (eL), nitrogen deficiency (N-) or combinations thereof. Four and seven days after treatment induction the expression of indicated genes was determined in four biological replicates per treatment. Results are expressed as log_2_(fold change compared to the control sample). Significance is indicated for p<0.001 as asteriks, or an FDR<0.01 for RNASeq data. *CHS - Chalcone synthetase, CHI - Chalcone isomerase, F3H - Flavanone-3-hydroxylase, FLS - Flavonol synthase, F3’H - Flavonoid-3’-hydroxylase, F3’5’H - Flavonoid-3’,5’-hydroxylase, DFR - Dihydroflavonol-4-reductase, LDOX - Leucoanthocyanidin dioxygenase*.

**Fig. 7:**
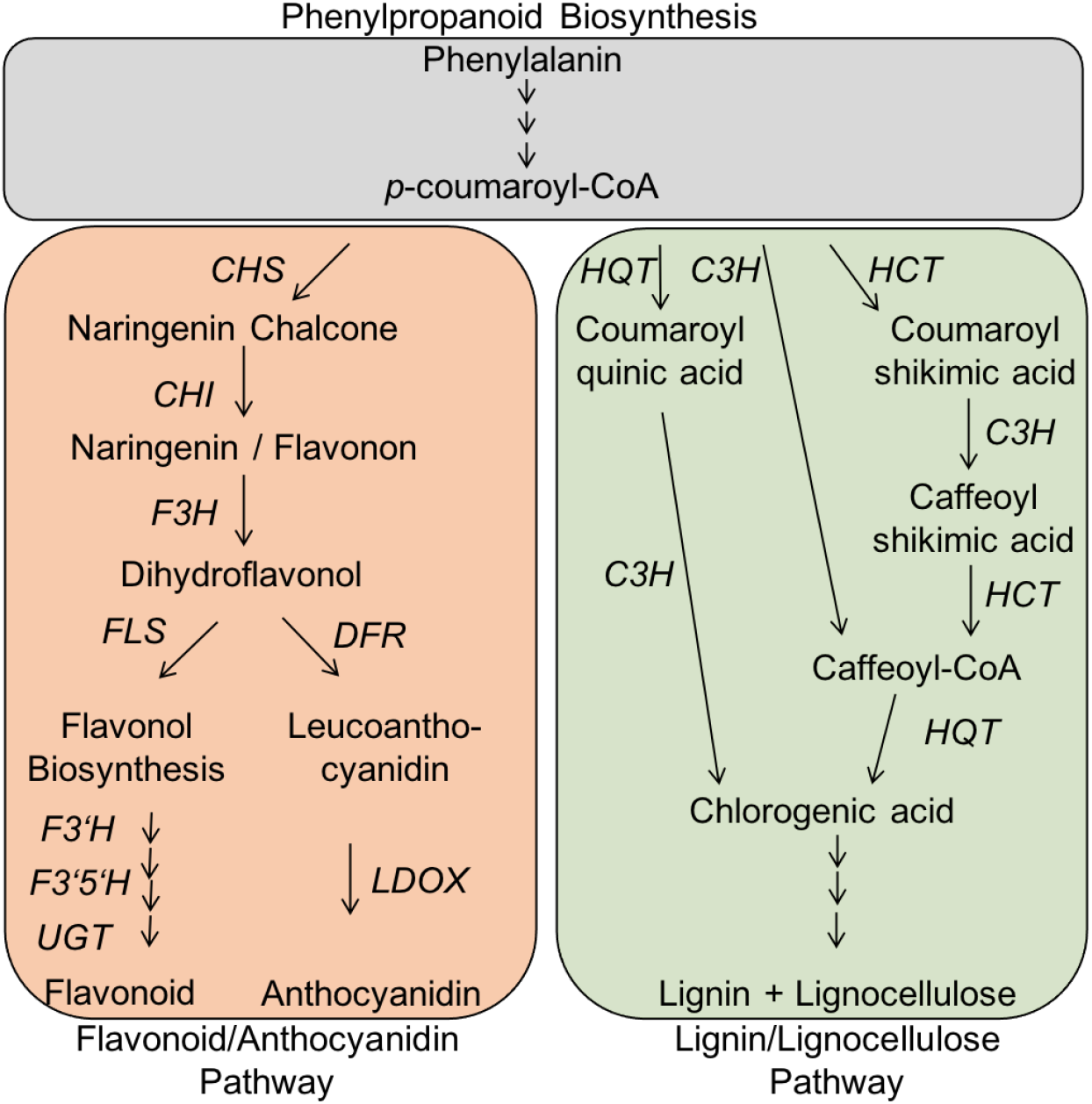
Comparative schematic overview of biosynthesis pathways of flavonoids, anthocyanidins and lignin and their structural genes. *p*-coumaroyl-CoA (grey box, top) is often the precursor for the flavonoids/anthocyanidin biosynthesis (red box, left side) or production of lignin or lignocellulose (green box, right side). Shown in this graphic are common pathways with their involved genes (adapted from André et al. (2009); Mahesh et al. (2007); Albert et al. (2009); Carvalho Lemos et al. (2019); Liu et al. (2020)). *CHS: Chalcone synthase, CHI: Chalcone isomerase, F3H: Flavanone 3-hydroxylase, FLS: Flavonol synthase, F3’H: Flavonoid 3’ hydroxylase, F3’5’H: Flavonoid 3’5’ hydroxylase, UGT: UDP-glycosyltransferase, DFR: Dihydroflavonol 4-reductase, LDOX: Leucoanthocyanidin dioxygenase, HQT: Hydroxycinnamoyl-Co A quinate hydroxycinnamoyl transferase, C3H: p-Coumarate 3-hydroxylase, and HCT: Hydroxycinnamoyl-Co A shikimate hydroxycinnamoyl transferase*.

In total, we found an up-regulation of structural genes involved in the flavonoid and anthocyanidin biosynthesis in both species four and seven days after stress treatment. In addition, the expression of structural genes is often higher induced after seven days when compared with day four. The induced expression of genes involved in the production of flavonoids is more pronounced for the cultivated tomato than in the wild relative (genes *FLS, F3’H* and *F3’5’H*). In contrast, genes involved in production of anthocyanidins are stronger induced in *S. pennellii* than in the cultivated tomato (genes *DFR* and *LDOX*). These findings support our observation of the metabolite analysis, where a larger quantity of flavonoids was detected for *S. lycopersicum* than for *S. pennellii* (see figure 4).

## 4 Discussion

Many environmental stresses, result in oxidative stress (Abewoy Fentik, 2017; Ksas et al., 2015; Chokshi et al., 2017; Robles-Rengel et al., 2019; Munjal and Munjal, 2019; Begara-Morales et al., 2019; García-Martí et al., 2019). As an often shown consequence, biosynthesis of plant secondary metabolites (PSM) is stimulated (Løvdal et al., 2010; Selmar and Kleinwächter, 2013; Grace, 2007; Dixon and Paiva, 1995) to overcome the stress (Wu et al., 2014; Isah, 2019; Aguirre-Becerra et al., 2021) and / or to become more tolerant (Nakabayashi et al., 2014). In our study we compared the effect of relatively mild abiotic stresses (warmer or chilling temperature regime, elevated light intensities or nitrogen deficiency, and combinations thereof) on the transcriptomic and metabolic response in a cultivated tomato (*S. lycopersicum*) with a well studied, stress tolerant wild relative (*S. pennellii*) in the vegetative state.

Overall, we found only mild induction and repression of pathways for elevated light intensities. Changed temperature regimes caused different behaviour in the cultivated variety and the wild relative. Nitrogen deficiency in any applied combination had the highest impact on transcriptional and metabolic response.

### Light response

Tomato is a photophilous crop, growing best under strong light intensities. In a previous study, a high light stress regime was described resulting in up to 120x higher light intensity than our control (24,000 µmol m^−2^ s^−1^ vs. 200 µmol m^−2^ s^−1^, respectively) (Parrine et al., 2018). Such a high light intensity exposure was accompanied by heat production up to 120 °C at the leaf surface. As a result damages in the photosystem pathways were described in combination with an oxidative stress response.

The applied light intensity in our experiment is equivalent to a cloudy (not overcast) midday in winter. Our elevated light regime almost doubled the light intensity, and simulates a shady location at midday illuminated by clear blue sky. As we did not measure the temperature of the leaf or within the canopy, we can only speculate about an additional warming due to radiation (see also discussion below). The applied increased light intensity caused only minor and mild effects on the transcriptome and metabolome in both species (see also figures 1, 4, 5 and supplementary figure 3).

As a result of the elevated light intensities, the investigated metabolites of the secondary metabolism (see figure 4) showed an induction up to 4-times in the cultivated tomato (see supplementary figure 10 a). Similar, induction of secondary metabolism by increased light intensities in the greenhouse was described before for *S. lycopersicum* (Løvdal et al., 2010; Junker-Frohn et al., 2019; Röhlen-Schmittgen et al., 2020). Interestingly, the wild relative *S. pennellii* did not respond to this stress treatment or repressed the production of the investigated secondary metabolites.

### Tomato’s response to changed temperature regimes

Although temperature plays an important role as a key regulator of plant growth and many further physiological processes (Hatfield and Prueger, 2015; Hildebrandt, 2018; Zhang et al., 2019), the applied mild changes in temperature regimes (warmer or chilling temperatures) in this study resulted in relative mild responses of the metabolome in both species.

Interestingly, the two species reacted different to the applied single stresses, while the transcriptome of the cultivated tomato responded stronger to warmer temperatures, chilling temperatures affected strongly the transcriptome of the wild relative (see figure 1 and supplementary figure 3). Those differences were as well visible for the investigated organic acids (see figure 5), but not for the investigated secondary metabolites (see figure 4).

Additionally, we found in the weighted cluster network analysis that samples under a single **warmer temperature regime** clustered separately in both species. Further, we were able to identify three related modules in the clustered network analysis for *S. lycopersicum* associated with the binary trait of warmer temperatures, but none for *S. pennellii* (see supplementary figure 6 and 7), suggesting that the wild species might be less responsive to the mild temperature stress.

For cultivated tomatoes a highly orchestrated temperature regime depending on the developmental stage is discussed (Boote et al., 2012; Shamshiri et al., 2018). It includes optimal temperature ranges of 18 – 21 °C during night and 21 – 30 °C during day for all developmental stages (Camejo et al., 2005; Shamshiri et al., 2018), and especially 22 °C as being optimal for the vegetative state (Sato et al., 2000; Adams et al., 2001), and 24 – 25 °C for optimized leaf area expansion, shoot elongation and plant biomass (Ntatsi et al., 2014), in addition to fruit development and maturation (Adams et al., 2001). In this regard, the applied warmer temperature regime was still in the optimal temperature range for tomato development.

Nevertheless, we observed the induction of small heat shock proteins in both species, which is a common effect of increasing temperatures (Lindquist, 1986; Lindquist and Craig, 1988). Two of these genes were also found in module brown for *S. lycopersicum* (Solyc01g102960.3.1 and Solyc11g020330.1.1).

In addition, adaptation to increased temperature can cause a photosynthetic response (Berry and Björkman, 1980), as we found it for *S. lycopersicum* (see supplementary figure 4). However, we did not see any changes in genes related to photosynthesis for *S. pennellii* (see supplementary figure 5), implying that photosynthesis is not affected in the stress tolerant wild species by our chosen temperature regime. In our data nitrogen deficiency has a higher impact resulting in a decreasing photosynthetic capacity (due to transcriptomic data) in both species, as we discuss later.

Further, it is described for tomato, that warmer temperature regimes can result in increased auxin and gibberellin signalling accompanied by cell wall reorganisation (Ohtaka et al., 2020). In our study the induction of auxin signaling pathways were observed in *S. lycopersicum*, but gibberellin signaling was repressed. Regarding cell wall reorganisation, an induction of arabinogalactan protein pathways as well as cuticular lipid formation was observed. In addition, cluster network analysis identified two expansin genes (Solyc06g060970.2.1 and Solyc08g077900.3.1) and a cellulose synthase like gene (Solyc12g088240.2) as being associated with the response to warm temperatures.

In contrast, cell wall reorganisation pathways (cellulose-hemicellulose network assembly, pectin modification, lignin polymerisation, cutin biosynthesis) were repressed in the wild species *S. pennellii*, but phytosterol biosynthesis was induced. It has been shown previously that plants increase the levels of sterol glucosides as a response to high temperature to maintain membrane integrity (Mishra et al., 2015; Narayanan et al., 2016).

Additionaly, the ORA revealed an induction of solute transport via the ATP binding cassette (ABC) transporter superfamily. In *S. lycopersicum*, that could indicate increased transport of cutin monomers across the plasmamembrane into the apoplast. With the help of the weighted cluster analysis we identified three additional genes belonging to the solute transport category: Solyc03g019820.3.1 and Solyc06g072130.4.1 are tonoplast intrisic proteins, that are involved in transport of water and small molecules across the vacuolar membrane, and Solyc06g051860.3.1 is a PHT1-type phosphate transporter. Aquaporins are known to play an important role in maintaining cell homeostasis under abiotic Stress (Afzal et al., 2016; Kapilan et al., 2018; Kurowska, 2021) and future studies may investigate their role in tomato’s response to warm temperature.

Our data also revealed that pathways of protein S-glutathionylation were repressed in both species, although for Arabidopsis a flavonoid-mediated induction of glutathione S-transferase12 (At5 g17220) was described (Nakabayashi et al., 2014). Usually S-glutathionylation of proteins is a reaction of oxidative stress, and glutathion is an important component to scavenge superoxide radicals, that might result from photosynthetic and respiratory electron transport chains (Dixon et al., 2005; Gurrieri et al., 2019; Dumont and Rivoal, 2019). Therefore, we assume that the applied warm temperature regime did not result in an oxidative stress.

The **chilling temperature regime** altered the expression pattern of genes in the wild species *S. pennellii* much stronger in contrast to the cultivated tomato. For *S. lycopersicum* we found only a small number of common genes up-regulated under either chilling temperatures or the combination of chilling temperatures and elevated light intensities, belonging to various biological functions.

Especially for *S. lycopersicum* we noticed, that genes within the identified module were either up-regulated under chilling temperature or a combination of chilling temperature and elevated light intensities, or down-regulated under any other stress combination. We focused on up-regulated genes under cold or eLcold, as we were mainly interested in induction of genes (see table 4). Here, five genes were identified, that were up-regulated under both stress treatments. For Solyc02g079370.3.1 (cyclin D 6;3), Solyc03g111950.3.1 (cytochrome P450), and Solyc12g055920.3.1 (Calcineurin B-like protein 4) further studies are necessary to uncover their role in response to chilling temperature. Solyc12g009190.2.1 (protein kinase (LRR-III)) is described to be involved in salt and drought stress response (Zhu et al., 2018).

The identified Solyc08g063080.3.1 is a putativ UDP-sulfoquinovose synthase. This group of proteins is involved in the biosynthesis of sulfolipid sulfoquinovosyl diacylglycerol. Sulfoquinovosyl diacylglycerol is a component of plant photosynthetic membranes and the most saturated glycolipid (Janero and Barrnett, 1981; Sanda et al., 2001; Brychkova et al., 2013). This class is the only sulphur-containing anionic glycerolipid, and maintains thylakoid fluidity, stabilization of the photosynthetic processes, in particular ATP synthesis, and photosystem II functioning (Taran et al., 2000; Dörmann and Hölzl, 2009; Kobayashi, 2016). Thereby induction of Solyc08g063080.3.1 may help in membrane adjustment to temperature as discussed before (Scotti-Campos et al., 2019).

For the wild species, photosynthesis pathways were induced and genes for perception of phytohormones become activated under the chilling temperature regime. This is in accordance with a former publication showing that chilling temperatures result in a photosynthetic response (Zhou et al., 2018), which indicates a higher stress tolerance (Allen and Ort, 2001; Cao et al., 2015). In this respect *S. pennelllii* is a valuable source for breeding programs to integrate cold-stress tolerance in cultivated tomatoes.

We identified as well some modules related with the binary trait of chilling temperature in the weighted cluster analysis (see also supplementary figure 8 and 9). Genes belonging to the module purple in *S. pennellii* were induced under all stresses harboring chilling temperatures. Here, we identified a sucrose-phosphate synthase, Sopen09g035000.1. These synthases are involved in sugar metabolism and known to play a key role in adaptation to cold temperature (Pollock and Lloyd, 1987; Jones et al., 1998; Allen and Ort, 2001; Sasaki et al., 2001; Bilska-Kos et al., 2020). Sopen03g022310.1 and Sopen08g003230.1 (both HD-ZIP I/II transcription factor) belong to a class of genes that are described to be involved in plant growth and fruit ripening (Lin et al., 2008; Harris et al., 2011; Shao et al., 2018; Gu et al., 2019). Sopen04g006840.1 belongs to the Fumarylacetoacetate (FAA) hydrolase family. This family regulates the light response (Han et al., 2013; Zhi et al., 2019) and has been shown to play a role in salt stress response (Huang et al., 2018).

Nevertheless most of the identified, and up-regulated genes related with chilling temperatures in our data are yet of unknown function (see table 4), and further studies may elucidate their role in the response to chilling temperatures.

### Double stress eLcold has different effects than the single stresses in *S. pennellii*

In contrast to the cultivated tomato, the transcriptome of *S. pennellii* did not show an additive effect for the combination of chilling temperature and elevated light intensities, but it seems that the combination of these stresses gives a less pronounced response.

Chilling temperature affects many processes such as photosynthesis, carbohydrate metabolism, polyamine synthesis, reactive oxygen species (ROS) scavenging, protein folding, stabilizing cell structure, cell membrane integrity, and chromatin remodelling (Janmohammadi et al., 2015; Banerjee et al., 2017; Kosová et al., 2018; Fürtauer et al., 2019; Friedrich et al., 2019; Ritonga and Chen, 2020). In addition, it is known that light will influence the acclimation to chilling and cold temperature (Wanner and Junttila, 1999; Soitamo et al., 2008).

In our experimental set up we elevated plants for 0.8 m to double the light intensities. This may result also in an increase in the canopy and leaf temperature due to radiation and warmth produced by the lamps. In previous studies, additional radiation in our range increased the canopy temperature by 0.38 - 1 K (Nelson and Bugbee, 2015; Van Westreenen et al., 2020). In this respect, chilling stress might have been less pronounced in the combination of elevated light intensities and chilling temperatures during day conditions. Nevertheless, night time temperatures would not be affected by this and would still represent a chilling stress. In addition, well watered plants, as in our set up, chill their leaves by opening the stomata for evaporative cooling (Thomas and Prasad, 2003; Urban et al., 2017; Li et al., 2018b), this will antagonize the aforementioned points and contribute to a higher quantity to the leaf temperature (Nelson and Bugbee, 2015). Therefore, we assume, that also under the combination of elevated light intensities and chilling temperatures the canopy temperature is still a bit cooler during day and 6 K cooler during night than under control condition.

Chilling temperatures induced photosynthesis pathways as well as pathways belonging to histone-mediated chromatin organisation in *S. pennellii* (see supplementary figure 5 and section 3.2) in contrast to a combination of chilling temperature and elevated light intensities. Here, the elevated light intensities might neutralize the response to chilling temperatures in these pathways, as it was reported before for warm-climate plants (Allen, 2000; Allen and Ort, 2001; Scott Holaday et al., 2016).

In addition, chilling temperatures orchestrated pathways of protein homeostasis. This reaction was not observed for elevated light intensities condition or a combinatorial stress. We conclude, that in a combinatorial stress such a response is not necessary anymore due to the cold-repressed photosynthetic and respiratory activity (Allen and Ort, 2001; Scott Holaday et al., 2016).

The combination of light intensities and chilling temperatures induced genes of the terpene biosynthesis in *S. pennellii*. Terpenes are a wide group of unsaturated hydrocarbons originating from isoprene. Recently, the family of terpene synthase genes in tomato was characterized (Zhou and Pichersky, 2020), pointing to their influence in photosynthesis, electron transport, developmental regulation and membrane architecture (Pichersky and Raguso, 2018). In our study, either mild and undetected responses to the single stresses had an additive effect resulting in a detectable outcome only in the stress combination, or the double stress may activate additional pathways resulting in the detected overrepresentation.

To summarize, our transcriptional data of the wild species *S. pennellii* indicate well that the combination of chilling temperature and elevated light intensities, will not necessarily result in an induced response, but can suspend reactions in distinct pathways.

In contrast, our data of the metabolic screen show an additive effect of the stresses (see figure 4, 5 and supplementary figure 10). While warmer temperature did not show any induction of the investigated metabolites, chilling temperatures resulted in changes up to 17-fold induction of CQA II in *S. lycopersicum* and 2.5-fold induction of caffeoylglucaric acid III in *S. pennellii*. The combination of elevated light intensities and chilling temperatures further boosted the induction up to 37-fold induction of CQA II in *S. lycopersicum* and and 6.5-fold induction of caffeoylglucaric acid III in *S. pennellii*. As we did not observe corresponding transcriptional changes in the ORA, we assume, that the detected induction of metabolites might be a result of accumulation.

### Nitrogen deficiency and secondary metabolism

In this study nitrogen deficiency was not a sudden event (see methods 2.2). When the nitrogen-free medium is applied, the nitrogen stock within the plant will be consumed over time. Therefore, nitrogen deficiency in our experimental set up is only a mild stress with a delayed onset.

The performed principal component analysis, the weighted cluster analysis as well as the metabolic analysis indicated for both species that nitrogen deficiency has a more prominent effect on the transcriptome and metabolome than any other applied stress, and will not be attenuated in combination with another stress (in particular visible in supplementary figure 1 b).

Plants need nitrogen in a balanced concentration to grow properly and to avoid any toxic effects (for the plant but as well the environment) (Crawford and Forde, 2002; Masclaux-Daubresse et al., 2010; Aczel, 2019; Britto and Kronzucker, 2002). On the other hand, insufficient nitrogen supply when applied during the vegetative state was shown to not affect the plant height and thereby the plant growth and yield of tomato in general (Han et al., 2014). Furthermore, only little effects are described for insufficient nitrogen supply (50 % reduction) applied during onset of fruit ripening, which were line-specific, and only slightly reducing the absolute amounts of fruits but increasing the relative amounts of marketable fruits, while chemical content, concentration of secondary metabolites, flavour and taste were not affected (Hartz and Bottoms, 2009; Zotarelli et al., 2009; Duan et al., 2019; Schmidt and Zinkernagel, 2021). Nevertheless, reduction of nitrogen supply to 15-35 % of optimal conditions may result in up to 30 % lower yields due to lower fruit weight (Wang et al., 2015; Qu et al., 2020). Therefore, application of nitrogen deficiency to induce PSM production in leaves, and extracting them for industrial use, might be performed after the last fruit harvest as proposed formerly (Junker-Frohn et al., 2019; Röhlen-Schmittgen et al., 2020).

It is known that nitrogen deficiency will influence genes involved in cell cycle organisation, photosynthetic response, metabolism, plant hormone signal transduction (e.g. abscisic acid, auxin, or jasmonate), transporter activity, and oxidative stress responses (Roggatz et al., 1999; de Groot et al., 2004; Menz et al., 2016; Hsieh et al., 2018; Halpern et al., 2019), and it is hypothesized, that photosynthesis and growth process might be impeded, which could lower the crop yield (Mu and Chen, 2021). Furthermore, nitrogen deficiency can stimulate remobilization of stored nitrogen and the release of ammonium via the biosynthesis of phenylpropanoids (Richard-Molard et al., 2008; Krapp et al., 2011).

Our analysis is in accordance with the aforementioned points. In particular, we found for both species a specific repression of pathways belonging to the phytohormone action, photosynthesis pathways and cell cycle organisation and an induction of pathways belonging to RNA biosynthesis, solute transport and the secondary metabolism (see supplementary figure 4 and 5), what may result in the production of PSM, as it was also shown before for *Arabidopsis thaliana* and rice (Krapp et al., 2011; Cai et al., 2012).

The induction of the solute transport might be an effect of the organ-specific experimental set-up. From *Arabidopsis thaliana* it is known, that the shoot of the plant will support the roots by remobilization of nitrogen as response to a sudden nitrogen deficiency (Krapp et al., 2011), and the same might be true in tomato.

However, we did not find a substantial number of transcription factors enriched in either the ORA or the weighted cluster analysis suggesting that involved transcription factors may have been more pronounced in the early phase of the answer, than in the present mid- to long-term response.

Many metabolic process were affected by the applied stresses harboring nitrogen deficiency in both species. And a few times, we observed an induction and repression of the same main category in the ORA, see as an example the behaviour of secondary metabolism under eLcold and N-cold for *S. lycopersicum* in table 2. This pattern is a result from different sub-categories indicating that different specific pathways within that main category are induced or repressed due to the abiotic stress treatment. In this context, biosynthesis of terpenes is repressed in *S. lycopersicum*, while biosynthesis of flavonoids is induced (see supplementary figure 4). Interestingly, this behaviour could not be observed for the wild species (see supplementary figure 5).

With the help of the weighted cluster analysis we were able to identify a set of up-regulated genes that might be related to the response to nitrogen deficiency in both species (see table 3). In the cultivated variety *S. lycopersicum* we identified Solyc05g014710.4.1, a putative remorin. Remorins are plant-specific and plasma membrane-associated proteins (Raffaele et al., 2007; Gui et al., 2016). They are discussed to be involved in responses to abiotic stress such as high salinity, chilling temperature or drought (Kreps et al., 2002; Malakshah et al., 2007; Jarsch and Ott, 2011; Checker and Khurana, 2013; Yue et al., 2014; Sowiński et al., 2020), and biotic stresses (Jarsch and Ott, 2011; Lefebvre et al., 2010; Raffaele et al., 2007; Gui et al., 2016). So far, remorins have not been describe in connection with nitrogen deficiency.

Sopen03g025580.1, also a candidate detected in the weighted cluster analysis, is a member of the Mlo family, in *S. pennellii*. Genes of the Mlo family are known to be important for the resistance against powdery mildew. In addition to resistance to powdery mildew, Mlo genes also participate in a variety of biotic and abiotic stress responses (Kim and Hwang, 2012; Kim et al., 2014; Lim and Lee, 2014; Nguyen et al., 2016; Acevedo-Garcia et al., 2017). For tomato it was shown that Mlo genes are involved in responsiveness to absisic acid, methyl jasmonate, heat stress and salicylic acid (Zheng et al., 2016). To our knowledge our observation is a first description that Sopen03g025580.1 is responding to abiotic stress caused by nitrogen deficiency.

In addition, we identified genes known to be involved in secondary metabolism such as: Solyc05g053550.3.1, a chalcone synthase, and Solyc10g052500.2.1, an isoflavone reductase homolog, in *S. lycopersicum*, and Sopen10g032910.1, a flavonol-3-O-rhamnosyltransferase, in *S. pennellii*. Among the further identified genes from the weighted cluster analyis, we found three putative uridine 5’-diphosphate (UDP)-glycosyltransferase (UGT) for *S. lycopersicum* (Solyc04g082860.3.1, Solyc10g085880.1.1 and Solyc04g074340.3.1), and two putative UGT for *S. pennellii* (Sopen11g003340.1 and Sopen03g021390.1).

Among all to date identified UGTs, only a few of them have been biologically characterised. *In vitro* studies have identified phenylpropanoids and flavonoids as UGT substrates, while their *in vivo* function remains elusive (Chong et al., 2002; Gachon et al., 2004; Kim et al., 2006; von Saint Paul et al., 2011; Langenbach et al., 2013; Simon et al., 2014; Song et al., 2015; Liu et al., 2018). In case of flavonoid biosynthesis, UGTs are involved in the last steps (see figure 7 left side). They have a conserved Plant Secondary Product Glycosyltransferase (PSPG) motif (Gachon et al., 2005), that we found as well in our identified UGTs by using expasy PROSITE (https://prosite.expasy.org). This motif is involved in the binding of the UDP moiety of the sugar molecule (Gachon et al., 2005). Glycosylation in general affects the toxicity, stability, complexity, spectral characteristics and solubility of flavonoids and other secondary metabolites (Vogt and Jones, 2000; Bowles et al., 2005). This is often essential for flavonoid transport, storage and signal transduction (Jones and Vogt, 2001).

Four of the investigated PSM were glycosylated derivates of either kaempferol or quercetin (see figure 4), with a higher abundance of quercetin derivates. Glycosylation is a common modification of secondary metabolites in plants (Wang and Hou, 2009). In general, glycosylated secondary metabolites have a reduced chemical activity compared to the aglycone, and therefore, they act merely as storage and/or a transport form (Li et al., 2001).

It was shown that chloroplasts can participate in the biosynthesis of flavonoids (Saito, 1974). Additionally, roots grown in complete darkness do not accumulate flavonoids due to light dependent expression of genes encoding flavonoid biosynthesis-related enzymes, as shown for *Arabidopsis thaliana* (Jenkins et al., 2001). As a consequence, leaves are discussed as the major tissue in which flavonoid biosynthesis occurs in higher plants (Deng et al., 2019). Additionally, flavonoids can be transported long distances from their synthetic sites to distant tissues (root tips) through the vascular system (Buer et al., 2007).

For the cultivated tomato we found an induced production of several PSM as seen in the performed metabolic analysis (see figure 4), with highest amounts found under the triple stress N-eLcold. This observation was accompanied by bins for the flavonoid synthesis enriched in up-regulated genes (see supplementary figure 4), while genes belonging to the biosynthesis of terpenoids were repressed. Induction and parallel repression of specific secondary metabolite biosynthesis pathways were described before in *Arabidopsis thaliana* (Scheible et al., 2004). In addition, the contrary behaviour of flavonoid and terpenoid biosynthesis depending on nitrogen availability was shown before for *Rosmarinus officinalis* (Bustamante et al., 2020).

Kaempferol and quercetin derivates are known to be induced under combinatorial stresses of nitrogen deficiency, chilling temperatures and / or elevated light intensities in cultivated tomato lines (Løvdal et al., 2010; Junker-Frohn et al., 2019; Röhlen-Schmittgen et al., 2020). Our observed induction rate is in the range of the previously mentioned publications. Interestingly, the wild relative *S. pennellii* showed much lower induction rates for the investigated metabolites (see supplementary figure 10). This is in contrast to a publication of Schauer et al. (2004). He indicated a higher content of secondary metabolites in leaves of *S. pennellii* compared to a cultivated variety by analysing the quinate and shikimate content. It might be that *S. pennellii* induced other PSM than the investigated metabolites in our study. This can result in the observed difference between our data and Schauer et al. (2004).

It is well published, that kaempferol and quercetin derivates assist in the response to abiotic and biotic stresses (Nakabayashi et al., 2014; Mierziak et al., 2014). But the dihydroxy B-ring-substituted quercetin derivates can interact as well with light, in contrast to the monohydroxy B-ring-substituted kaempferol derivates (Agati et al., 2012; Fini et al., 2011; Mierziak et al., 2014). Besides, production of quercetin derivates can be induced by light and nitrogen deficiency (Stewart et al., 2001; Kanazawa et al., 2012; Xie et al., 2012), while kaempferol production will be up-regulated under fertilisation (Deng et al., 2019). This might be a reason, for the observed ratio between quercetin and kaempferol derivates. In addition, the higher accumulation of quercetin derivates might result in a higher drought and oxidative tolerance as it was shown before for Arabidopsis (Nakabayashi et al., 2014).

Besides several PSM belonging to the flavonoid synthesis pathway, we found a strong induction of three regioisomers of mono-caffeoylquinic acid in both species (see figure 4). With our experimental approach we were unable to further elucidate the structure of the mono-caffeoylquinic acid regioisomers, but it is highly probable that the identified regioisomer II is equivalent to chlorogenic acid. Chlorogenic acid (5-O-caffeoylquinic acid) is a member of the group of mono-caffeoylquinic acids, and belongs to the biosynthesis of lignin (part of the cell wall). Both biosynthetic pathways, either resulting in the production of flavonoids and anthocyanidins or lignin and lignocellulose, start from p-coumaroyl-CoA (see figure 7). Caffeoylglucaric acid, the last identified PSM, is known to be synthesized in tomato from chlorogenic acid and glucaric acid by chlorogenic acid:glucaric acid caffeoyltransferase (Strack et al., 1987; Strack and Gross, 1990).

Chlorogenic acid is not only a central intermediate in the biosynthesis of lignin. It can be found in a wide variety of foods and beverages, including fruits, vegetables, olive oil, spices, wine, and especially coffee (Pérez-Jiménez et al., 2010; Socała et al., 2021), as well as tobacco leaves (Sheen, 1973). Along with chlorogenic acid (5-O-caffeoylquinic acid, also called 5-CQA), 4-O-caffeoylquinic acid (also called neochlorogenic acid) and 3-O-caffeoylquinic acid (also called cryptochlorogenic acid) appear as well in the plant kingdom (reviewed in Liu et al. (2020)), while 1-O-caffeoylquinic acid was seldom isolated (Parejo et al., 2004). 4-O-caffeoylquinic acid is identified in various edible and medicinal plants, such as mulberry (*Morus alba L.*), *Acanthopanax henryi*, and *Ilex kudingcha* (Ganzon et al., 2018; Zhang et al., 2014; Xu et al., 2015). Furthermore, chlorogenic acid has been identified by an untargeted metabolome analysis in tomatoes resistant for *Alternaria alternata*, as the main inhibitor of the alternariol biosynthesis (Wojciechowska et al., 2014).

Caffeoylquinic acids have been investigated in different diseases because of its anti-oxidative, antibacterial and antiparasitic effect, anti-inflammatory activity, reduction of chronic pain, such as neuroprotective, anticancer, cardiovascular and antidiabetic effects (Scholz et al., 1994; Li et al., 2018a; Santana-Gálvez et al., 2017; Bagdas et al., 2019; Socała et al., 2021). In particular, caffeoylquinic acids were investigated for the treatment of cancer, cardiovascular disease, diabetes, cognitive impairment, hypercholesterolemia, nutritional and metabolic diseases, and impaired glucose tolerance (reviewed in Liu et al. (2020)).

In addition to the induction of PSM, we observed a reduction of produced metabolites belonging to the primary metabolism (see figure 5) in both species. In particular malic acid, aconitic acid and citric acid were reduced under nitrogen deficiency, while quinic acid, *β*-D-gluconic acid and D-glucoheptonic acid became induced.

Malic acid, aconitic acid and citric acid are intermediates of the citric acid cycle. Quinic acid is known to be a main part in the biosynthesis of caffeoylquinic acids, such as chlorogenic acid (Clifford et al., 2017), and is widely occuring in the plant kingdom. It is discussed, that primary and secondary metabolism may compete for the available photosynthetic assimilates (Stamp, 2004). We conclude, that especially the cultivated tomato *S. lycopersicum* curbs the primary metabolism to respond to the nitrogen deficiency stress by induction of the secondary metabolism, as it was shown before for *Arabidopsis thaliana* (Scheible et al., 2004).

## 5 Conclusions

Our study is a first detailed transcriptomic and metabolomic analysis of response to various mild, abiotic stresses and combinations thereof in leaves for the cultivated tomato variety *S. lycopersicum* and its wild relative *S. pennellii*. We focused on a mid- to long-term timepoint to get insights on the underlying molecular mechanisms. Nevertheless our analysis is only a snapshot of one timepoint and one specific organ, that is rarely investigated. Further studies will be necessary to investigate and differentiate mid- and long-term response in different tomato organs (e. g. root, shoot, different ages of leaves, flower,…).

We were able to show substantial alterations on the molecular level based on our transcriptome and metabolome analysis as a response to the stress treatment but also between the species. This study shows that leaf transcriptome and metabolite levels are under great environmental influence, as mild abiotic stresses result in strong transcriptomic and metabolic responses.

In our experimental set-up nitrogen deficiency had the highest effect in both species, and both induced widely the phenylpropanoid metabolism. But the effect was more pronounced in the leaves of the cultivated tomato, resulting in higher accumulation of PSM and a suppression of the primary metabolism, while the wild species only slightly repressed the primary metabolism. Overall our study indicates leaves of tomatoes under a mild abiotic stress as a valuable source for antioxidants such as flavonoids or mono-caffeoylquinic acid. In addition, our analysis indicate a more diverge response in the wild species, reflecting as well a breeding potential of *S. pennellii* to generate e. g. a cold-tolerant cultivated tomato.

## Availability of data and material

The following material is available online: Supplementary figure 1-10, Supplementary Table S1: Table of DEG, Supplementary Table S2: Table of metabolites. RNASeq data is available at the NCBI: BioProject PRJNA641520.

## Code availability

Not applicable

## Author contributions

Conceptualization: Julia J. Reimer, Laura V. Junker-Frohn, Anika Wiese-Klinkenberg and Alexandra Wormit; RNASeq analysis: Julia J. Reimer; Metabolomic analysis: Björn Thiele; validation: Julia J. Reimer and Robin T. Biermann; formal analysis: Julia J. Reimer; writing–original draft preparation: Julia J. Reimer; writing–review and editing: all authors; visualization: Julia J. Reimer and Robin T. Biermann; project administration: Alexandra Wormit; funding acquisition: Anika Wiese-Klinkenberg and Alexandra Wormit.

