## Supplemental figures for "Tomato leaves under stress: A comparison of stress response to mild abiotic stress between a cultivated and a wild tomato species"

Journal: Plant Molecular Biology

Corresponding author: A. Wormit
(RWTH Aachen University
)

### Supplementary Figure 1

### PCA from a combined analysis of *S. lycopersicum* and *S. pennellii*

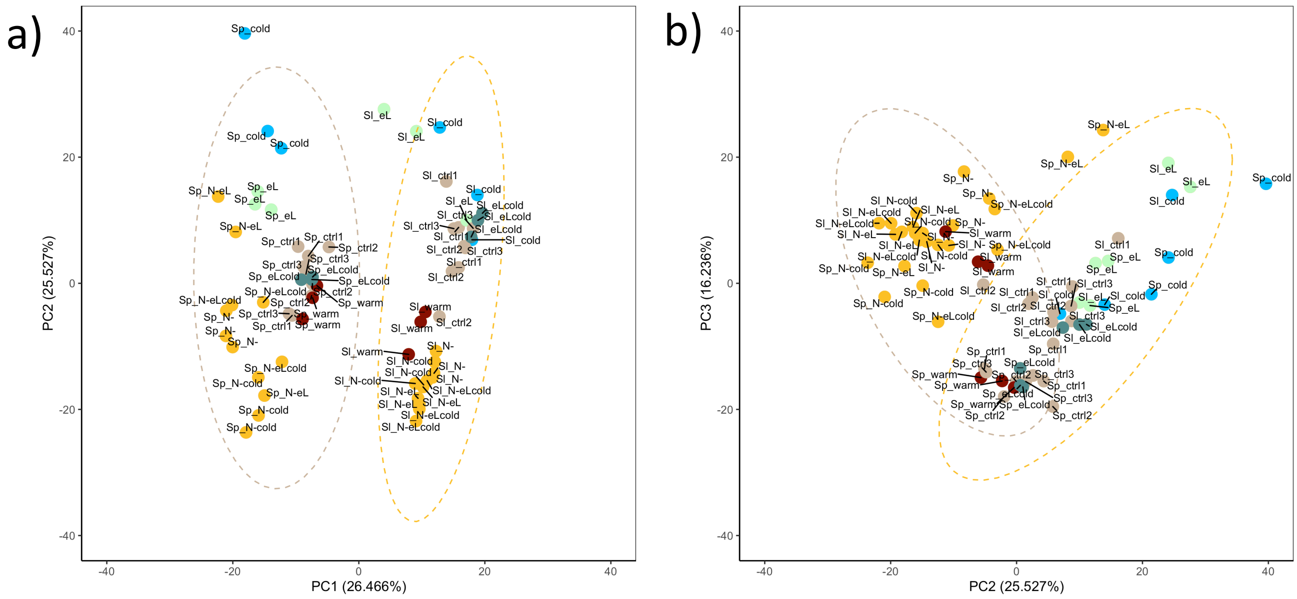


RNASeq reads were mapped to an artificial transcriptome comprising both transcriptomes coming from *S. lycopersicum* (Sl) and *S. pennellii* (Sp) (see also methods). Shown in a) is the principal component 1 (PC1) versus principal component 2 (PC2), indicating that the highest variation in the combined analysis is coming from the species (Sl versus Sp). In b), the principal component 2 (PC2) is shown over principal component 3 (PC3), indicating that the next level of variation is influenced by the stress treatments. Control samples are indicated in brown, cold samples in blue, warm samples in red, elevated light samples in light green, and all samples comprising nitrogen deficiency in orange.

Supplementary Figure 2

Dendrogram of trait cluster for *S. lycopersicum* and *S. pennellii*

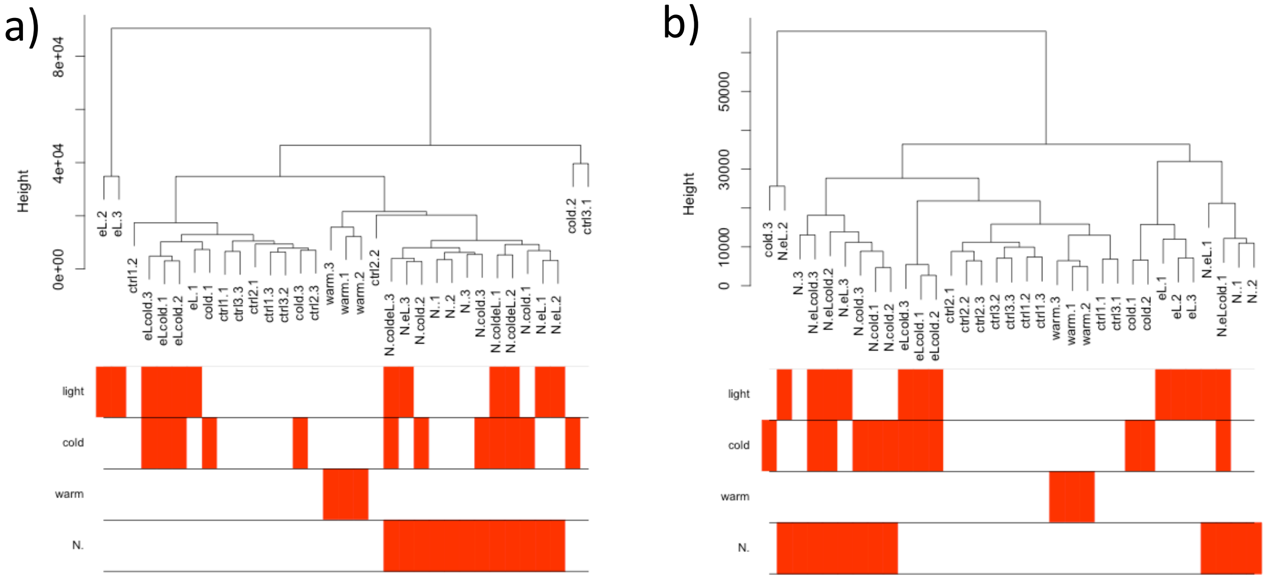


RNASeq reads were used to perform a cluster network analysis (see also methods) for *S. lycopersicum* (a) and *S. pennellii* (b). The dendrogramm shows what clusters will be formed according to the read abundance in combination with the single stresses. Stresses are indicated as N (nitrogen ), eL (elevated light intensities), cold (chilling temperature), warm (warmer temperature) or ctrl (control). Combinatory stresses are indicated as eLcold, N.cold, N.eL and N.coldeL. The number behind the stress indication reflects the single RNASeq sample.

Supplementary Figure 3

Volcano plots indicating the differentially expressed genes in *S. lycopersicum* (a) and *S. pennellii* (b)

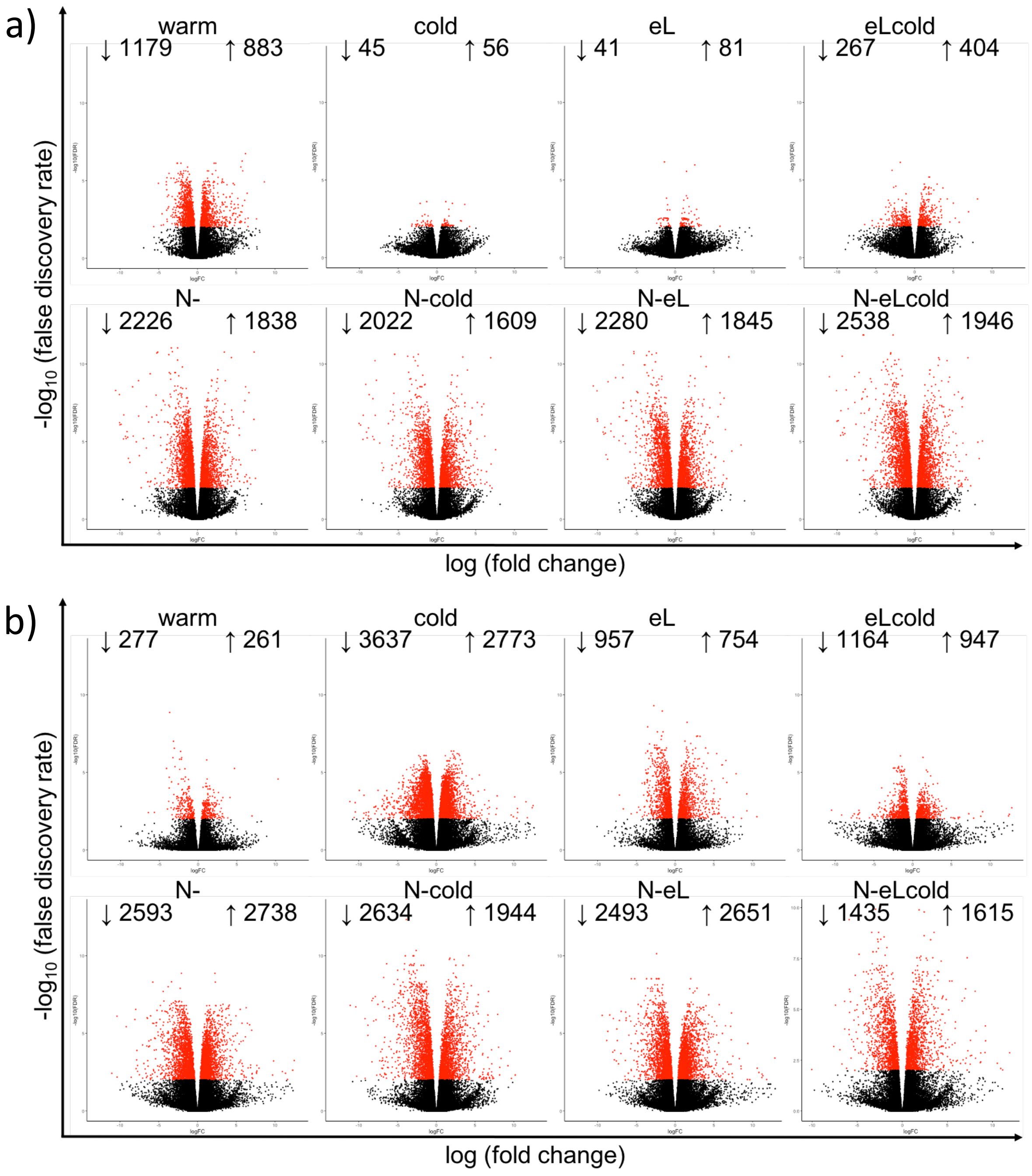


Plants were grown under different abiotic stress conditions (elevated light intensities: eL, chilling temperatures: cold, warmer temperature regime: warm, nitrogen deficiency: N-, and combinations thereof as indicated). The negative decadic logarithmic values ($-\log_{10}$) of the false discovery rate are ploted against the logarithmic values of the fold change for each condition in both species. Significantly different expressed genes are highlighted in red, and total numbers are indicated. FDR < 0.01

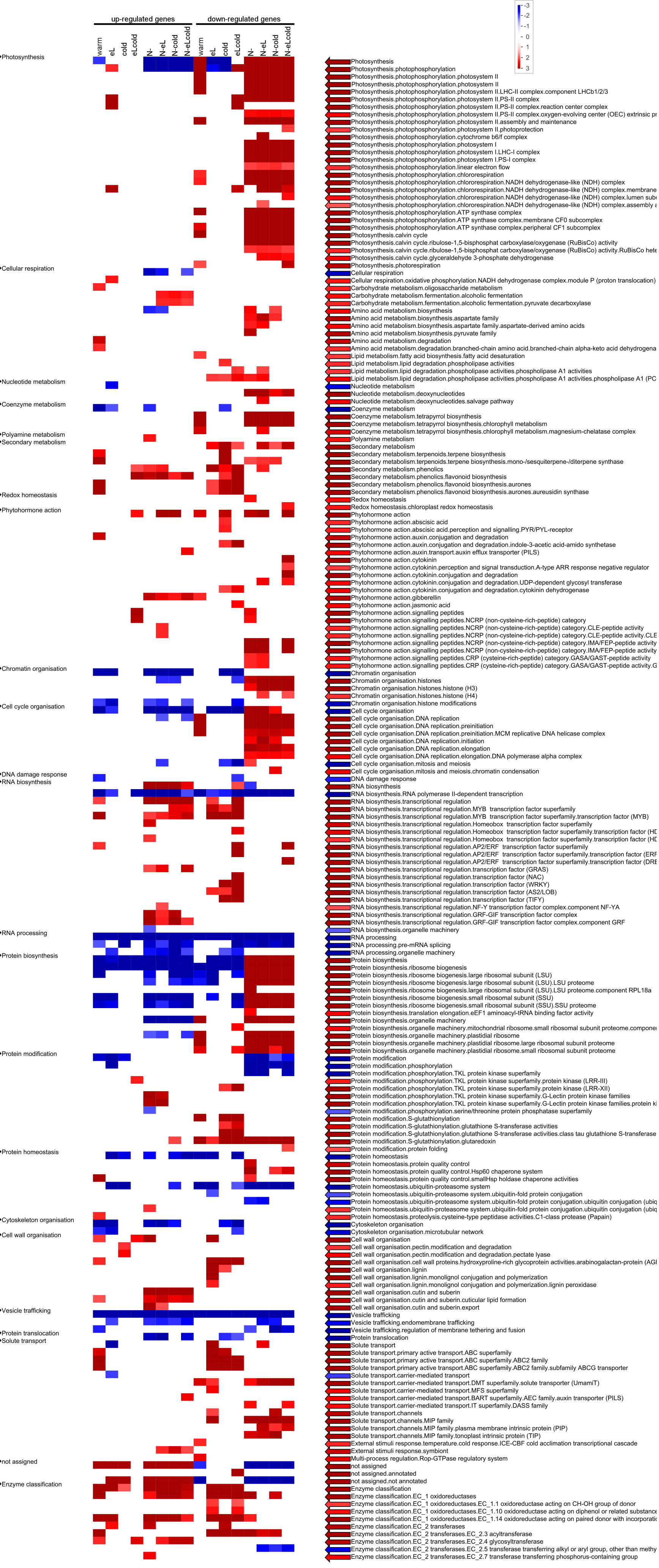


### Supplementary Figure 4 Overrepresentation analysis (ORA) performed for up- or down-regulated DEG of *S. lycopersicum*

The over- or under-representation of indicated Mercator bins (Schwacke et al., 2019) were calculated seperately for up- (left column) and down-regulated genes (right column) using a Fisher test (Usadel et al., 2006) in combination with a multiple testing correction as performed by Benjamini and Hochberg (Benjamini and Hochberg, 1995). Overrepresented bins are shown in red and underrepresented bins in blue, respectively.

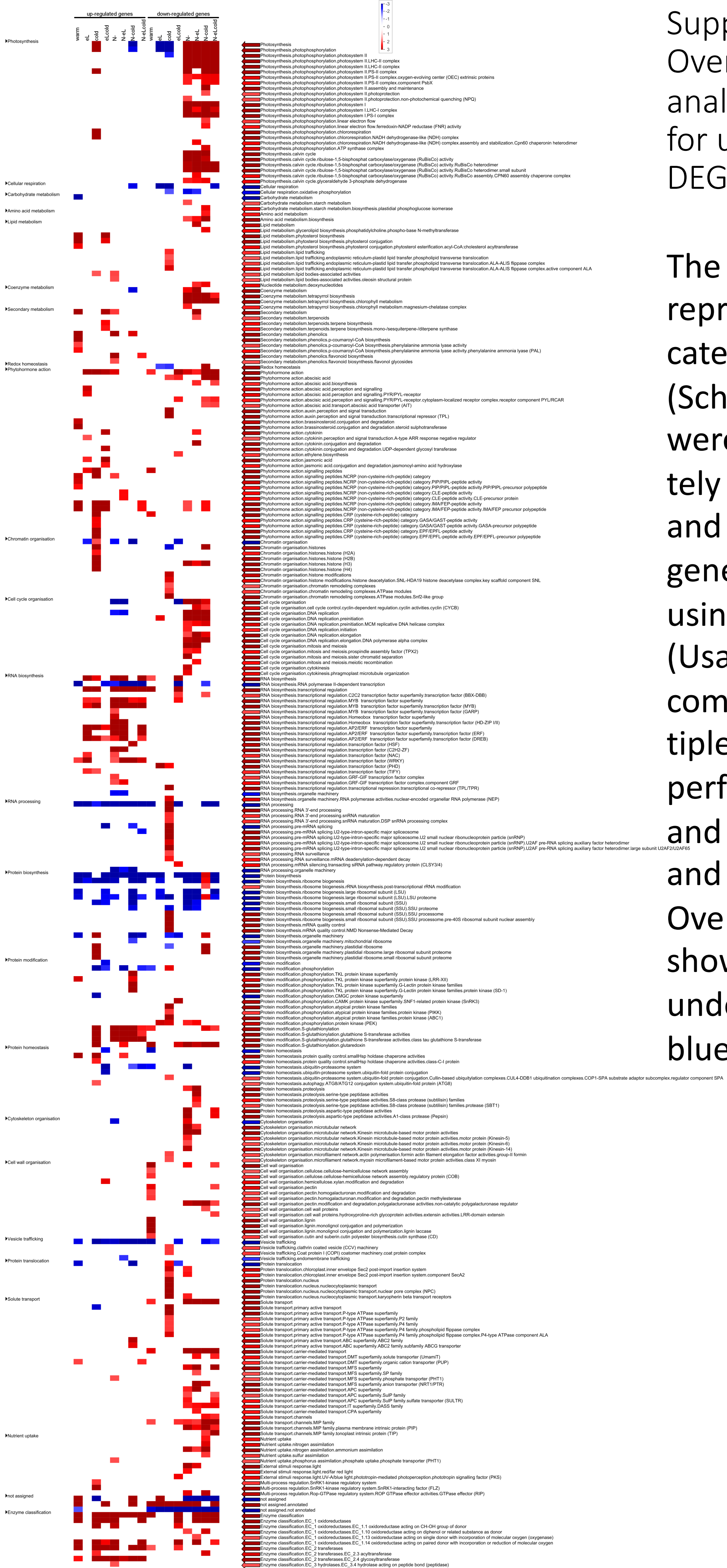

Supplementary Figure 5 Overrepresentation analysis (ORA) performed for up- or down-regulated DEG of *S. pennellii*

The over- or under-representation of indicated Mercator bins (Schwacke et al., 2019) were calculated seperately for up- (left column) and down-regulated genes (right column) using a Fisher test (Usadel et al., 2006) in combination with a multiple testing correction as performed by Benjamini and Hochberg (Benjamini and Hochberg, 1995). Overrepresented bins are shown in red and underrepresented bins in blue, respectively.

### Supplementary Figure 6
### Module-trait correlation for *S. lycopersicum*

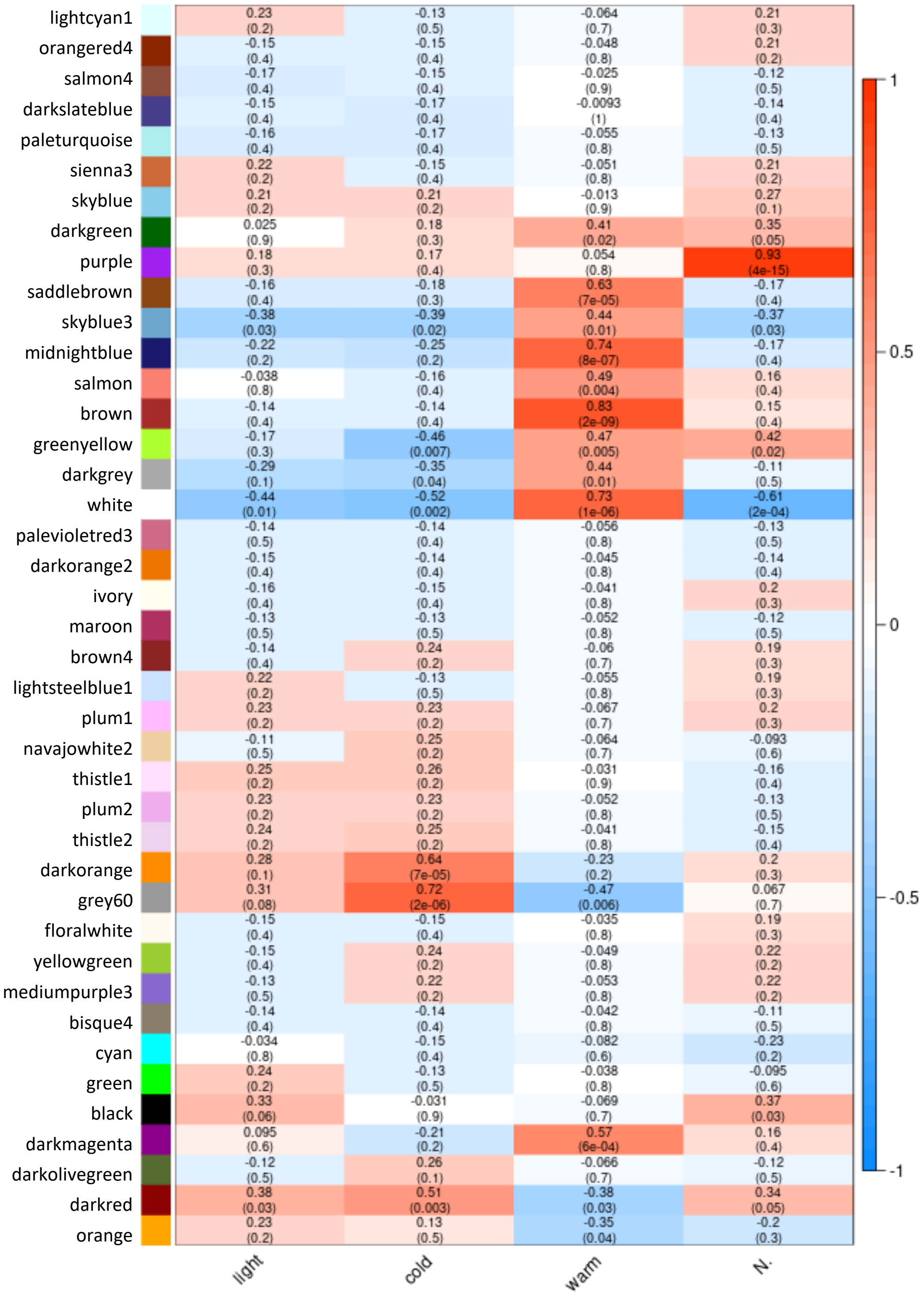


RNASeq data was used to perform a weighted cluster analysis. Shown here is the relationship for each identified cluster with the single stress elevated light intensities (light), chilling temperature (cold), warmer temperature (warm), or nitrogen deficiency (N) as indicated. On the y-axis, all identified modules are listed with there respective names. Color and number indicate the correlation between module and binary trait for eigengenes, and numbers in brackets the p-Value for significance.

Supplementary Figure 7
Module-trait correlation for *S. pennellii*

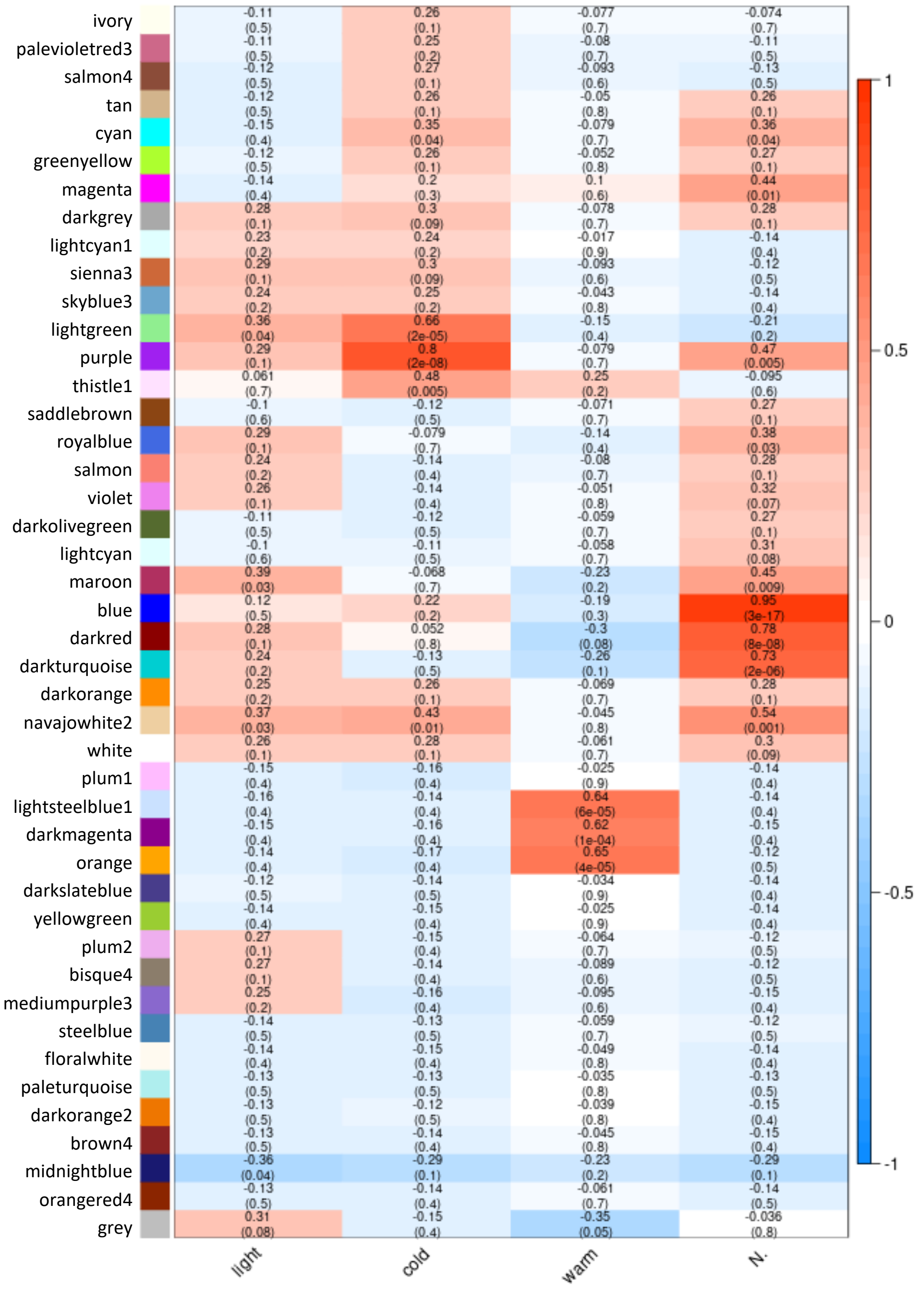


RNASeq data was used to perform a weighted cluster analysis. Shown here is the relationship for each identified cluster with the single stress elevated light intensities (light), chilling temperature (cold), warmer temperature (warm), or nitrogen deficiency (N) as indicated. On the y-axis, all identified modules are listed with there respective names. Color and number indicate the correlation between module and binary trait for eigengenes, and numbers in brackets the p-Value for significance.

Supplementary Figure 8
Characteristics of modules with a high correlation for one binary trait shown for *S. lycopersicum*

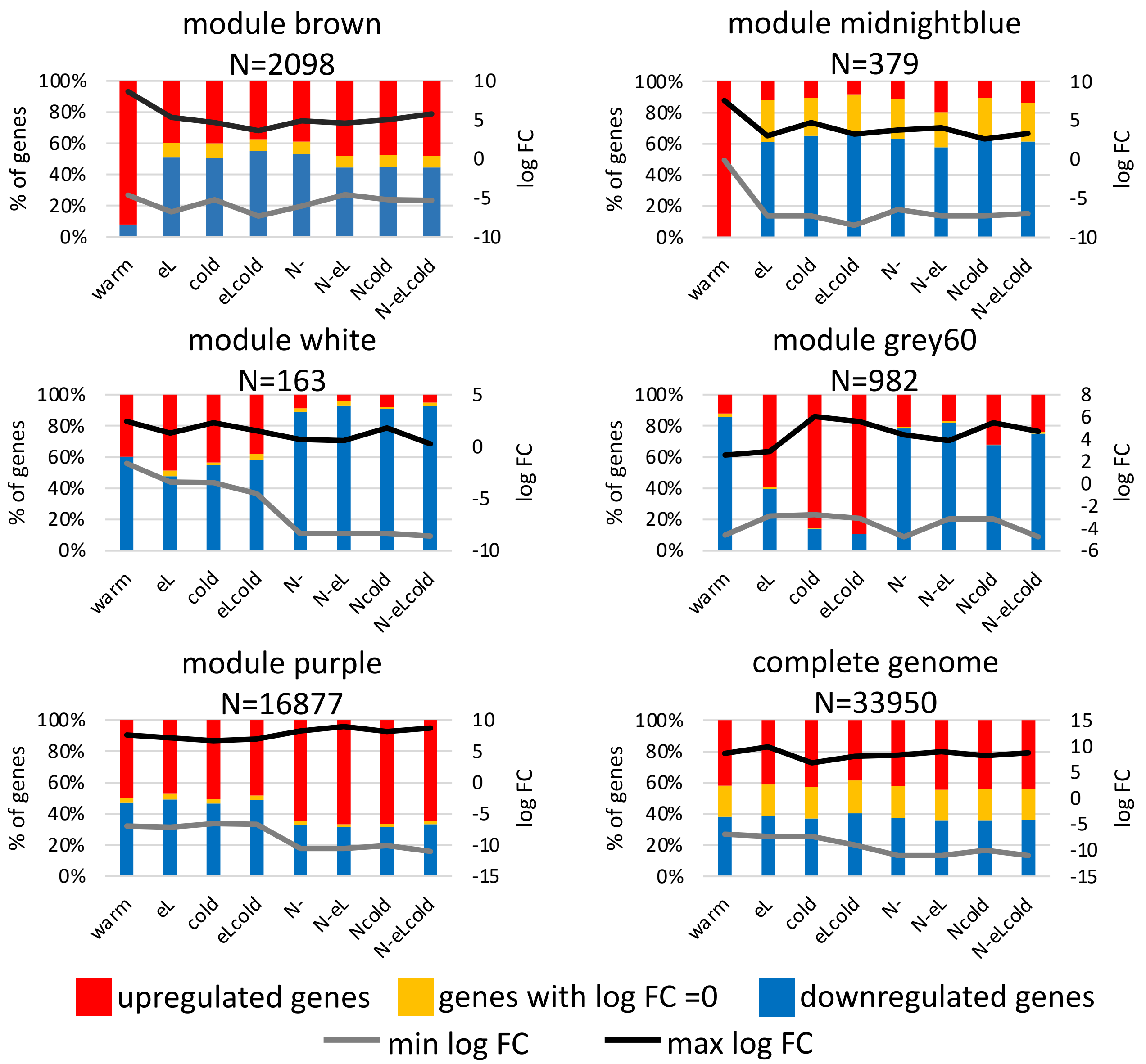


RNASeq data was used to perform a weighted cluster analysis. Modules with a correlation of 0.72 and higher were investigated for the presence of up- (red) or down-regulated (blue) genes. Genes having a $\log_2$ FC = 0 are indicated in yellow. The absolute maximal $\log_2$ FC (black line) and minimal $\log_2$ FC (grey line) are indicated. Beneath the module name is shown the number of genes within the module. The graph on the bottom right shows data for the whole genome.

Supplementary Figure 9

Characteristics of modules with a high correlation for one binary trait shown for *S. lycopersicum*

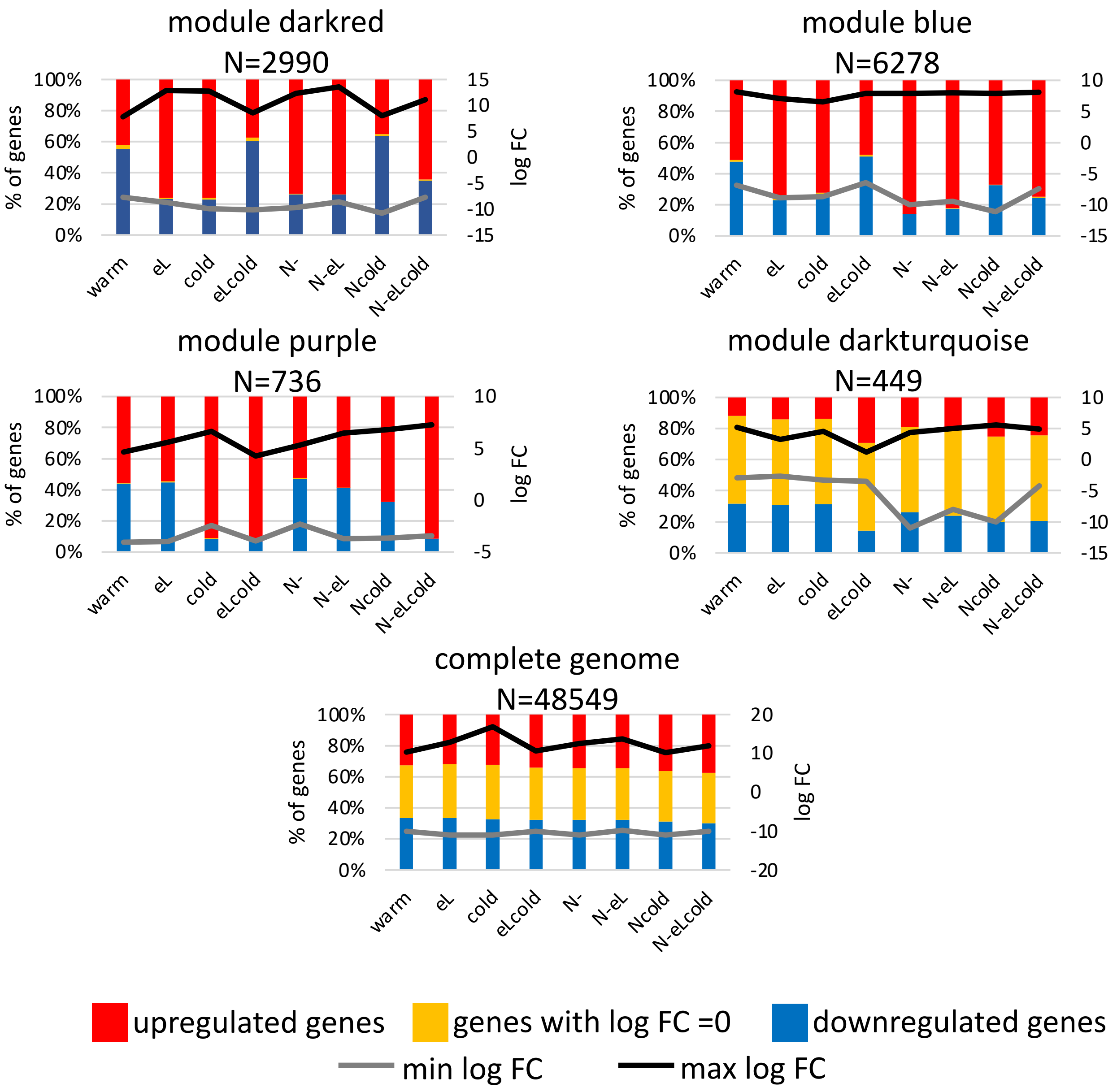


RNASeq data was used to perform a weighted cluster analysis. Modules with a correlation of 0.72 and higher were investigated for the presence of up- (red) or down-regulated (blue) genes. Genes having a $\log_2$ FC = 0 are indicated in yellow. The absolute maximal $\log_2$ FC (black line) and minimal $\log_2$ FC (grey line) are indicated. Beneath the module name is shown the number of genes within the module. The graph on the bottom shows data for the whole genome.

Supplementary Figure 10
Fold induction of detected metabolites in both species

a)

| | sample | CQA I | CQA II | CQA III | Caffeoyl glucaric acid I | Caffeoyl glucaric acid II | Caffeoyl glucaric acid III | Caffeoyl glucaric acid IV | Quer-3-O-Hex-DHex-Pent | Kaemp-3-O-Hex-DHex I | Kaemp-3-O-Hex-DHex II | Quer-3-O-Hex-Dhex II (Rutin) | Quer-3-O-Hex-Dhex I |
|---|---|---|---|---|---|---|---|---|---|---|---|---|---|
| *S. lycopersicum* | warm | 0 | -0.3 | -0.3 | -0.1 | -0.2 | -0.3 | 0.3 | 0.3 | 7.3 | -7.1 | -2 | 9.5 |
| | eL | 2.1 | 2.4 | 2.6 | 2 | 2.5 | 1.5 | 0.8 | 1 | 10.4 | -5.8 | -1.2 | 10.6 |
| | cold | 1.8 | 3.3 | 4.1 | 2.6 | 3.3 | 2.8 | 2.6 | 1.8 | 12.4 | -4.4 | -1 | 11.1 |
| | eLcold | 2.3 | 4.1 | 5.2 | 2.5 | 3.3 | 4.3 | 3.8 | 2.3 | 10.8 | -2.9 | -0.4 | 11.5 |
| | N- | 3 | 4.9 | 5.6 | 3.2 | 2.9 | 5.2 | 4.2 | 3.2 | 11.6 | -1.7 | 0.8 | 12 |
| | N-cold | 3.4 | 5.6 | 6.2 | 3.3 | 3.7 | 6.5 | 4.9 | 4.1 | 12.4 | -0.1 | 1.2 | 13 |
| | N-eL | 2.9 | 5.4 | 6.8 | 3.4 | 3.2 | 6 | 5.3 | 4.9 | 12.7 | -0.9 | 1.9 | 13 |
| | N-eLcold | 3.3 | 5.8 | 6.5 | 3.5 | 3.3 | 6.9 | 5.1 | 4.9 | 13.2 | 0.4 | 2 | 13.9 |
| *S. pennellii* | warm | n.d. | 0.1 | 0.9 | 0 | 0.3 | 0.3 | 1.1 | 0.6 | 4.7 | -4.5 | -0.4 | 6.5 |
| | eL | n.d. | 0.1 | 0.5 | -0.9 | -0.8 | -0.1 | 0.5 | -0.3 | 3.6 | -3.9 | -1.6 | 4.9 |
| | cold | n.d. | 0.9 | 1.2 | -1 | -0.1 | 1.3 | 1.2 | 0.2 | 3.8 | -2.5 | -1.2 | 5.4 |
| | eLcold | n.d. | 1.6 | 2.4 | 0.3 | 0.7 | 2.7 | 2.7 | 0.8 | 4.5 | -1.4 | -1.4 | 5.2 |
| | N- | n.d. | 1.6 | 2.3 | 0.4 | 1.1 | 2.4 | 2.4 | 1.3 | 4.9 | -1.4 | -0.4 | 5.9 |
| | N-cold | n.d. | 2.3 | 2.4 | 0.4 | 0.6 | 3.4 | 2.7 | 1.7 | 5.3 | 0.2 | -1.1 | 5.6 |
| | N-eL | n.d. | 1.8 | 3.7 | 1.5 | 1.8 | 3.2 | 3.9 | 3.8 | 7.4 | -0.4 | 0 | 7.4 |
| | N-eLcold | n.d. | 2 | 2.2 | 0.7 | 0.6 | 3 | 2.4 | 2 | 5.4 | -0.2 | -1.1 | 5.5 |

b)

| | sample | Malic acid | Aconitic acid | Citric acid | Quinic acid | β-D-Gluconic aicd | D-Glucoheptonic acid |
|---|---|---|---|---|---|---|---|
| *S. lycopersicum* | warm | 0.1 | 0 | 0 | -0.4 | 0.1 | 0.2 |
| | eL | 0 | 0 | -0.2 | 1.6 | 0.5 | 0 |
| | cold | 0 | -0.1 | -0.2 | 2.5 | 1.3 | -0.2 |
| | eLcold | 0 | -0.1 | -0.3 | 3.5 | 1.1 | -0.1 |
| | N- | -3.2 | -0.5 | -2.8 | 4.2 | 0.4 | 0.9 |
| | N-cold | -2.5 | -0.6 | -2.9 | 5.1 | 0.7 | 0.8 |
| | N-eL | -3.5 | -0.4 | -3.5 | 4.8 | 0.1 | 1 |
| | N-eLcold | -2.7 | -0.7 | -2.9 | 5.4 | 0.3 | 0.7 |
| *S. pennellii* | warm | 0.5 | 0.2 | 0 | -0.1 | 0 | 0 |
| | eL | 0.4 | 0 | -0.1 | 0.4 | 0.4 | 0.1 |
| | cold | 0.5 | 0 | 0.2 | 1.1 | 0.9 | 0.3 |
| | eLcold | 0.4 | -0.1 | 0.1 | 1.9 | 0.9 | 0.3 |
| | N- | -1 | -0.1 | -0.5 | 2 | 1.8 | 0.5 |
| | N-cold | -0.8 | -0.4 | -0.4 | 2.8 | 1.3 | 0.3 |
| | N-eL | -0.7 | 0 | -0.4 | 2.3 | 2.3 | 0.7 |
| | N-eLcold | -0.3 | -0.4 | -0.4 | 2.5 | 1.3 | 0.2 |

| magnitude of change | 0 | 1 | 2 | 3 | 4 | 5 | 6 | 7 | 8 | 9 | >10 |
|---|---|---|---|---|---|---|---|---|---|---|---|
| greater than zero | | | | | | | | | | | |
| less than zero | | | | | | | | | | | |

Samples used for the RNASeq analysis and the qPCR verification were used for determination of metabolites by LC-ESI(+)-FTICR-MS and either LC-ESI(+)-MS/MS (a) or LC-ESI(+)-MS/MS (b). All metabolites were unambiguously identified. Quantification of them was performed with LC-MS/MS by use of their corresponding MRM transitions. Results are expressed as $\log_2((\text{peak area ratio})_{treatment}/(\text{peak area ratio})_{control})$. The color code indicate $\log_2$ values in both tables. n.d. = not detected
